# Identification of the mode of evolution in incomplete carbonate successions

**DOI:** 10.1101/2023.12.18.572098

**Authors:** Niklas Hohmann, Joël R Koelewijn, Peter Burgess, Emilia Jarochowska

## Abstract

**Background:** The fossil record provides the unique opportunity to observe evolution over millions of years, but is known to be incomplete. While incompleteness varies spatially and is hard to estimate for empirical sections, computer simulations of geological processes can be used to examine the effects of the incompleteness *in silico*. We combine simulations of different modes of evolution (stasis, (un)biased random walks) with deposition of carbonate platforms strata to examine how well the mode of evolution can be recovered from fossil time series, and how test results vary between different positions in the carbonate platform and multiple stratigraphic architectures generated by different sea level curves.

**Results:** Stratigraphic architecture and position along an onshore-offshore gradient has only a small influence on the mode of evolution recovered by statistical tests. For simulations of random walks, support for the correct mode decreases with time series length. Visual examination of trait evolution in lineages shows that rather than stratigraphic incompleteness, maximum hiatus duration determines how much fossil time series differ from the original evolutionary process. Gradual directional evolution is more susceptible to stratigraphic effects, turning it into punctuated evolution. In contrast, stasis remains unaffected.

**Conclusions:**

- Fossil time series favor the recognition of both stasis and complex, punctuated modes of evolution.
- Not stratigraphic incompleteness, but the presence of rare, prolonged gaps has the largest effect on trait evolution. This suggests that incomplete sections with regular hiatus frequency and durations can potentially preserve evolutionary history without major biases. Understanding external controls on stratigraphic architectures such as sea level fluctuations is crucial for distinguishing between stratigraphic effects and genuine evolutionary process.

## Introduction

### The fossil record as source of information

Fossils provide a unique record of evolution on temporal and spatial scales not accessible to experimentation or direct human observation (Gingerich 1983; 2001). Geological records have delivered fossil time series crucial in formulating and testing hypotheses on evolutionary dynamics and mechanisms of speciation spanning micro-to macroevolutionary scales (e.g., Dzik 2005; Strömberg 2006; Aze et al. 2011; Voje 2020). Nevertheless, fossils remain underused in evolutionary biology. Their main application is still node and tip calibration of molecular clocks, which commonly relies on single occurrences or on assumptions about the probability of finding a fossil rather than stratigraphic data (Donoghue and Yang 2016). It is also subject to biases resulting from this small sample size (Springer 1995). The unique type of information contained in a fossil succession sampled over a long time interval is rarely exploited, likely due to the following barriers:

1. The fossil record, being a part of the stratigraphic record, is patchy and distorted. At the time when Darwin (1859) discussed this as a major limitation for the testing and development of the theory of evolution, little geological knowledge was present to elucidate the rules governing this incompleteness. Darwin’s concern widely persists (e.g., Patterson (1981)), albeit mostly implicitly: most phylogenetic analyses published today do not use fossils which would have been relevant or use only a small fraction of them. Stratigraphy and sedimentology, which can provide relevant data on fossils and the constraints on their occurrence and sampling (Kidwell and Holland 2002; Hunt 2010), are jargon-laden, highly atomized disciplines whose utility for evolutionary biology is not obvious to biologists. Biostratigraphy, which uses fossils to establish the relative age of rocks and has amassed datasets that would be of high utility for evolutionary studies, employs taxonomic concepts that are often impractical for or incompatible with evolutionary questions (Dzik 1985; 1995; Pearson 1992; Haug and Haug 2017). As a results, scientific communities studying evolution and the fossil record function in parallel, with limited exchange (Grantham 2004).
2. There is a lack of methodological frameworks to incorporate fossils in their stratigraphic context in evolutionary studies. Historically, phylogenetic methods rarely incorporated geological information such as the relative order of appearance of taxa or specimens in the fossil record, which is the main subject of biostratigraphy (Wills 1999). This has led to radical discrepancies between the outcomes of phylogenetic and stratigraphic, or stratophenetic, approaches (Gingerich and Schoeninger 1977; Donoghue 2001; Dzik 2005). This barrier is gradually overcome by methodological advances, such as the Fossilized Birth-Death Model (Stadler et al. 2018), which allows incorporation of parameters specific to the fossil record, such as fossilization rate, sampling probability and age uncertainties of fossil occurrences (Barido-Sottani et al. 2020; Warnock, Heath, and Stadler 2020; Wright et al. 2022; Barido-Sottani et al. 2023).

Recently, there is renewed appreciation for the importance of fossils in phylogenetic reconstructions (Quental and Marshall 2010; Mitchell, Etienne, and Rabosky 2019; Mongiardino Koch, Garwood, and Parry 2021; Guenser et al. 2021). These studies focus on the role of the morphological information provided by extinct taxa, but less on what a modern understanding of the physical structure of the geological record contributes to reconstructing evolutionary processes from fossil-bearing stratigraphic successions.

### Stratigraphic incompleteness and age-depth models

The incompleteness of the fossil record serves as an umbrella term for different effects that diminish the information content of the rock record, ranging from taphonomic effects and sampling biases to the role of gaps and erosion (Kidwell and Holland 2002). Here we focus on the role of gaps (hiatuses) in the rock record. Such gaps can arise due to sedimentation (including fossils) and subsequent erosion or lack of creation of rocks in the first place, e.g. when an environment remains barren of sediment formation or supply for a long time. Both processes result in gaps in the rock record and, as a result, in the fossil record. This type of incompleteness is termed stratigraphic (in)completeness, defined as the time (not) recorded in a section, divided by the total duration of the section (Dingus and Sadler 1982; Tipper 1987). Stratigraphic completeness provides an upper limit on the proportion of evolutionary history that can be recovered from a specified section, even with unlimited resources and perfect preservation of fossils. Stratigraphic completeness is difficult to quantify in geological sections, and estimates range between 3 and 30 % (Wilkinson, Opdyke, and Algeo 1991), suggesting that more than 70 % of evolutionary history is either not recorded in the first place or destroyed at a later time.

Fossils older than a 1.5 million years cannot be dated directly, and their age has to be inferred from circumstantial evidence on the age of the strata in which they were found (Wehmiller et al. 1988; Kidwell and Flessa 1996). This inference is formalized by age-depth models (ADMs), which serve as coordinate transformations between the stratigraphic domain, where the fossils were found (length dimension L, SI unit meter), and the time domain (time dimension T, SI unit seconds - we use the derived units years, kyrs, or Myrs) (Hohmann 2021b). Age-depth models are always explicitly or implicitly used when fossil data is used for evolutionary inferences. Because they convey how positions of fossils relate to their age, ADMs are the basis for calculating evolutionary rates. As a result, revising ADMs commonly leads to a revision of evolutionary rates. For example, Malmgren, Berggren, and Lohmann (1983) observed increased rates of morphological evolution in lineages of fossil foraminifera over a geologically short time interval of 0.6 Myr and proposed that this “punctuated gradualism” may be a “common norm for evolution”. MacLeod (1991) revised the age-depth model and showed that the interval with increased rates of evolution coincides with a stratigraphically condensed interval, i.e. more change is recorded in a thinner rock unit. Re-evaluating the evolutionary history based on the revised age-depth model removed the apparent punctuation and showed that morphological evolution in that case had been gradual rather than punctuated.

Age-depth models contain information on both variations in sediment accumulation rate and gaps in the stratigraphic and - as a result – the fossil record. For example, stratigraphic completeness corresponds to the fraction of the time domain to which an age-depth model assigns a stratigraphic position. In the absence of an age-depth model, we can only make statements on the ordering of evolutionary events, but not on the temporal rates involved.

### Forward models of stratigraphic architectures

Forward computer simulations of sedimentary strata provide a useful tool to study the effects of incompleteness and heterogeneous stratigraphic architectures. They have demonstrated that locations and frequency of gaps in the stratigraphic record are not random, but a predictable result of external controls, such as fluctuations in eustatic sea level (Warrlich 2000; Hutton and Syvitski 2008; Burgess 2013; Masiero et al. 2020).

Combined with biological models, forward models provide a powerful tool to test hypotheses on the effects of stratigraphic architectures on our interpretations of evolutionary history. For example, Hannisdal (2006) combined simulations of a siliciclastic basin with models of taphonomy and phenotypic evolution. The results showed that when sample sizes are small, morphological evolution will appear as stasis regardless of the underlying mode. This might explain why stasis is the most common evolutionary pattern recovered from the fossil record (Hunt, Hopkins, and Lidgard 2015).

Stratigraphic incompleteness and variations in sediment accumulation rates introduce multiple methodological challenges. Constructing complex ADMs requires sedimentological and stratigraphic expert knowledge, and they will potentially be associated with large uncertainties. Even in the “perfect knowledge” scenario where the age-depth model is fully known, evolutionary history in the time domain will inevitably be sampled irregularly: If two samples are separated by a hiatus, their age difference must be at least the duration of the hiatus, which might be millions of years. On the other hand, if sediment accumulation is rapid and no hiatuses are present, the age difference between samples might be only a few days.

Most studies “translate” fossil successions into time series using age-depth models based on simplified assumptions on the regularity of the stratigraphic record. These ADMs ignore stratigraphic incompleteness and often assume uninterrupted constant sediment accumulation (UCSA). This assumption implies that stratigraphic completeness is 100 %, rock thickness is proportional to time passed, and linear interpolation between tie points of known age can be used to infer fossil ages from their positions. Such ADMs are usually used implicitly, without discussing their limitations. While the assumption of UCSA is sedimentologically and stratigraphically unrealistic, it brings strong methodological simplifications. For example, if distance between samples collected in a rock section is kept constant, UCSA implies that the underlying evolutionary history in the time domain is sampled at a constant frequency, the generated fossil time series are equidistant in time and can therefore be analyzed by standard methods of time series analysis (Hunt 2006; Beran 2017).

### Objectives and hypotheses

We examine how commonly made simplified assumptions on stratigraphic architectures influence how the mode of evolution is recovered from fossil time series. We use tropical carbonate platforms as a case study, because they host large parts of the fossil record and are evolutionary hotspots (Jablonski, Roy, and Valentine 2006).

We test the following hypotheses:

1. The mode of evolution identified in a fossil time series obtained under the assumption of uninterrupted constant sediment accumulation (UCSA) is the same as the mode of the original time series.
2. Lower stratigraphic completeness reduces the chance of identifying the correct mode of evolution from fossil time series constructed based on the assumption of UCSA (Holland 2000; Patzkowsky and Holland 2012). The implication of this hypothesis is that different depositional environments have different chances of preserving the mode of evolution because of systematic differences in their completeness.

## Methods

We simulate trait evolution in the time domain, pass it through a stratigraphic filter produced by the CarboCAT Lite model of carbonate platform formation (Burgess 2013; Burgess 2023), and compare how well the mode of evolution can be recovered from the fossil time series sampled in the stratigraphic domain and the time series reflecting the “true” evolutionary history in the time domain. A visual summary of the workflow is shown in Figure 1. Data is available in (Hohmann et al. 2023), code is available (Hohmann et al. 2024). See the README and REPRODUCEME files therein for details on computational reproducibility.

**Figure 1:**
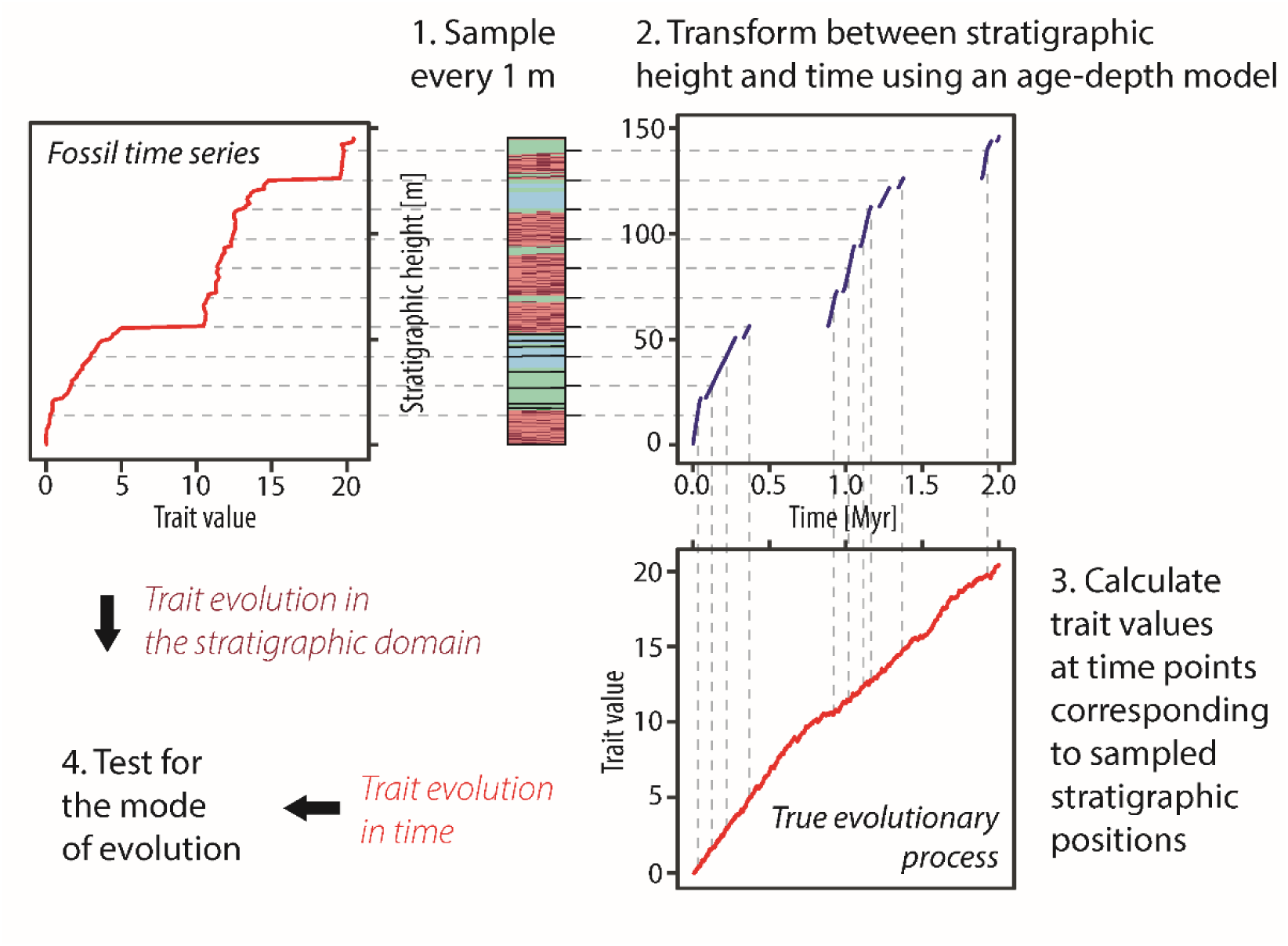
Study design for testing the mode of evolution in the stratigraphic domain. Computationally, first sampling positions are determined, then the age-depth model is used to determine the times that correspond to these positions. Last, the trait evolution at said times are simulated. The simulated mean trait values are the values observable at the sampled stratigraphic positions.

### The stratigraphic record: Forward models of carbonate platform architecture

We simulated two attached carbonate platforms using the CarboCAT Lite software (*Figure 2* and *Figure 3*) (Burgess 2023). CarboCAT Lite is implemented in MATLAB and simulates carbonate production by carbonate factories characterized by different production curves, which are functions of water depth (Bosscher and Schlager 1992; Burgess 2013; Masiero et al. 2020). It includes sediment transport that is a function of platform topography, i.e., sediment is transported downslope, but not a function of external parameters such as waves or currents. Simulations were run using time steps of 1 kyr (1000 years) and a grid of 1 km width (strike) and 15 km length (dip), subdivided into quadratic grid cells of 100 m length. We used three carbonate factories with production curves following the parametrization of Bosscher and Schlager (1992) (*Supplementary Figure 1*). Factory 1 is phototrophic with a maximum carbonate production rate of 500 m × Myr^-1^, which it maintains up to 30 m of water depth. Factories 2 and 3 have maximum production rates of 160 and 150 m × Myr^-1^ respectively and maintain maximum productivity until 40 m water depth, but differ in how fast productivity decreases with depth (*Supplementary Figure 1*). We assumed constant subsidence of 70 m per Myr across the grid. This is approximately seven times higher than estimated subsidence rates in the last 125 kyr in the Bahamas (McNeill 2005), allowing for higher completeness than would be generated on a typical passive continental margin. As initial topography, a slope with a gradient of 5.33 m × km^-1^ was used, resulting in a total initial difference in water depth of 80 m between the shore and the offshore end of the simulated grid. We followed the default setting of CarboCAT Lite, described by Burgess (2013) with three lithofacies (sediment types) corresponding to the three carbonate factories, plus three secondary lithofacies representing sediment transported downslope after the deposition of the primary lithofacies (dark colors in *Figures 2* and *3*). As initial facies distribution, the default setting of CarboCAT Lite was used, which assigns each grid cell one of the three carbonate factories or no factory according to a uniform distribution.

**Figure 2:**
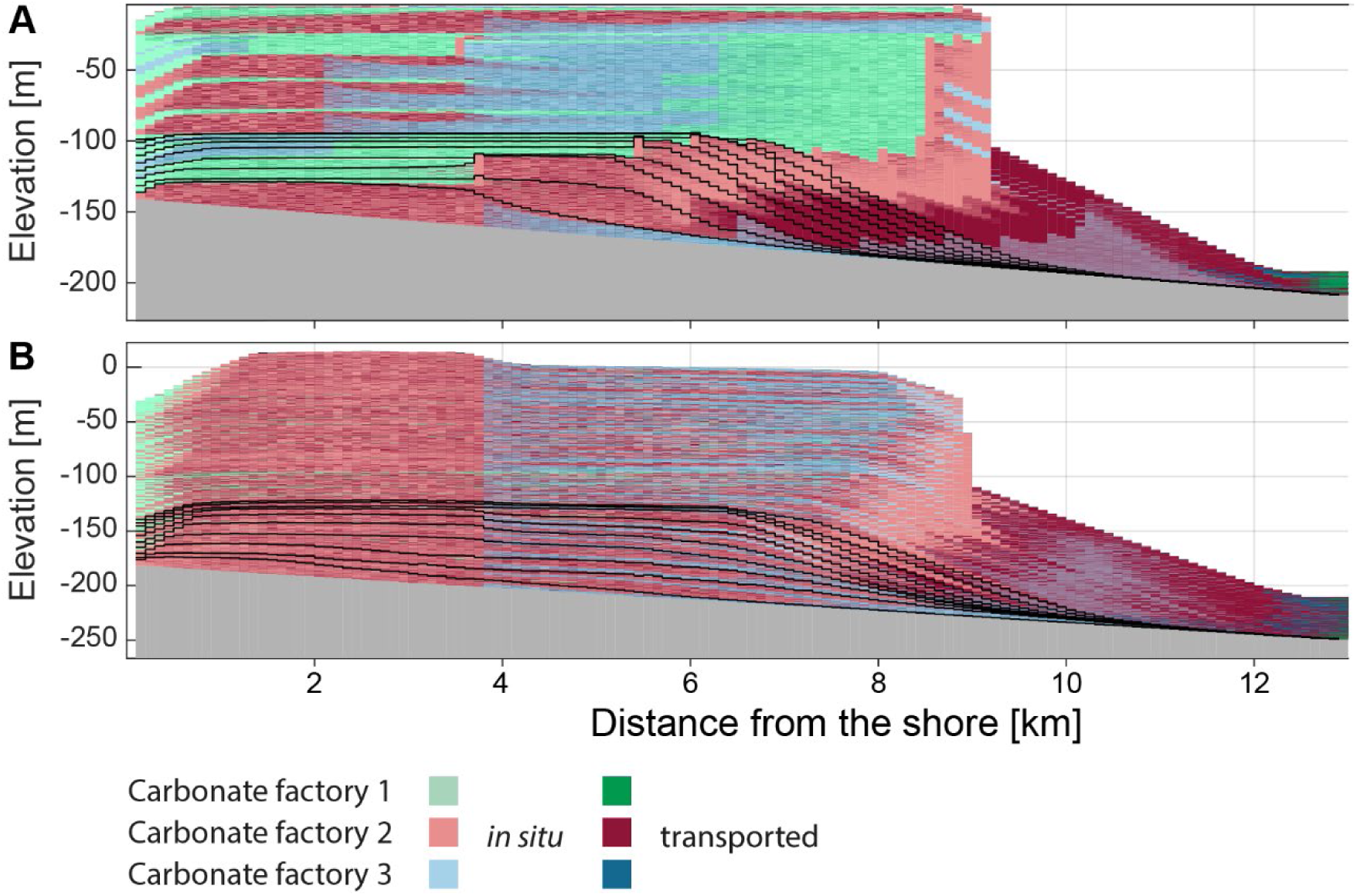
The outcome of simulating carbonate platforms in the stratigraphic domain. (A) Scenario A: deposition based on a fictional sea-level curve. (B) Scenario B: deposition based on the sea-level curve from Miller et al. (2020) for the last 2.58 Myr. Graphs represent the position in the middle of the simulated grid along the strike.

**Figure 3:**
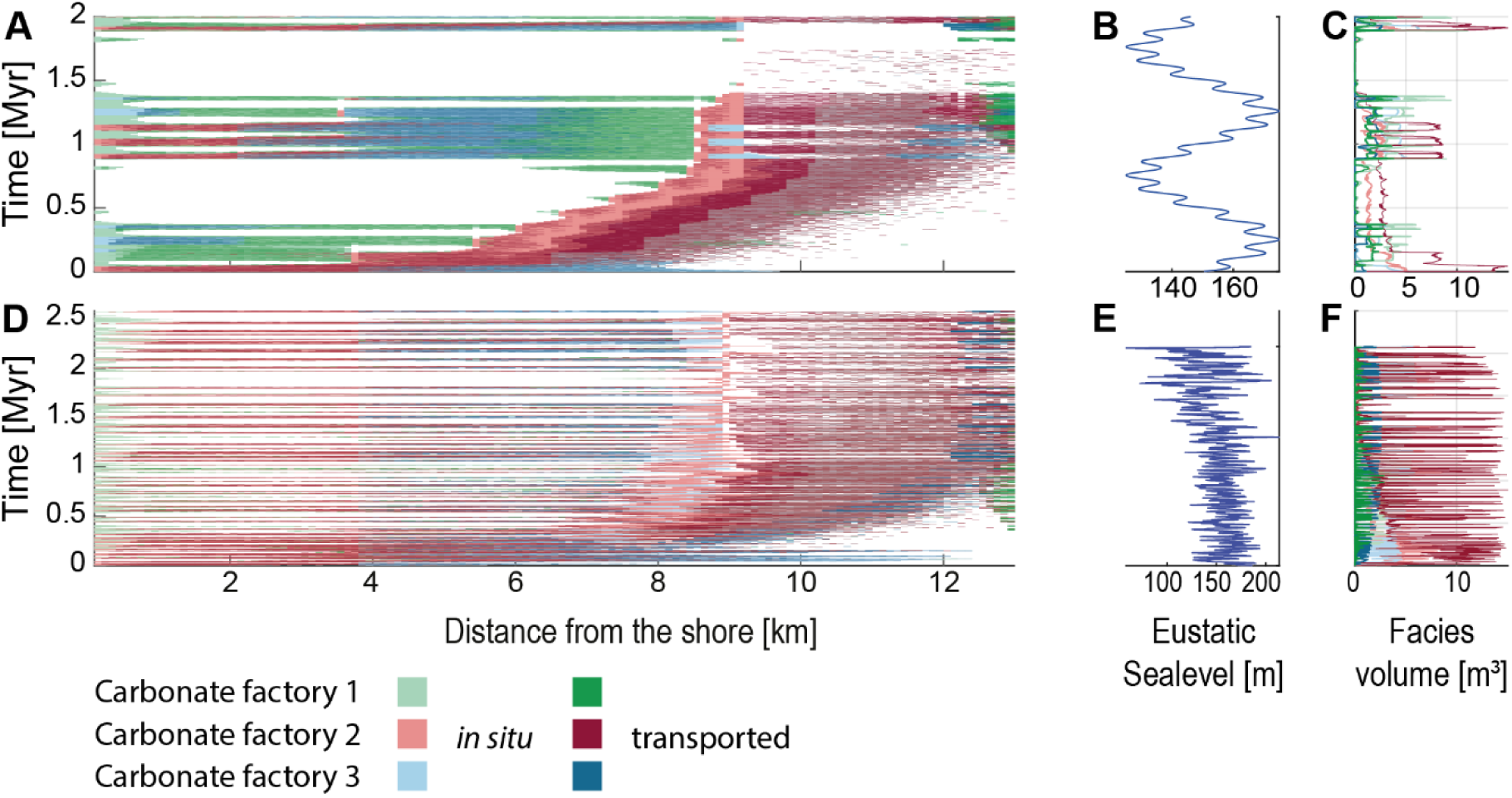
Simulated carbonate platforms in the time domain. (A-C) Scenario A. (D-F) Scenario B. (A, D) Chronostratigraphic (Wheeler) diagrams. (B, E) Sea level curves used as input for the simulation. (C, F) Facies volumes. Graphs represent the position in the middle of the simulated grid along the strike.

Two scenarios were simulated (*Figures 2* and *3*). In scenario A, the simulation was run for 2 Myr with changes in eustatic sea level given by a combination of sinusoids expressed as third-order sea level changes with a period of 1 Myr and an amplitude of 20 m and fourth-order changes with a period of 0.112 Myr and an amplitude of 2 m (*Figure 3 B*). Scenario B was simulated for the period from 2.58 Mya (beginning of the Pleistocene) until the present using the global eustatic sea level curve estimated from global δ^18^O_benthic_ records by Miller et al. (2020). This curve has a temporal resolution of approximately 2 kyr over the Pleistocene (mean: 2.13 kyr, median: 2.07 kyr), and was linearly interpolated to match the time increments of the simulation and normalized to have a mean sea level of 0 (*Figure 3 E*).

For each scenario, CarboCAT Lite produces a grid containing the facies, including no deposition, at each time step and position on the grid. The outputs were extracted from MATLAB and all further analysis was carried out in the R Software (R Core Team 2023). For both scenarios, age-depth models at every grid node were extracted from the simulation outputs. Age-depth models taken from the same distance from shore display only little variation as position along the shore (strike) is varied, indicating that lateral position in the grid is negligible. Accordingly, we focused on age-depth models taken from the middle transect along the dip. In each scenario, synthetic sections and the corresponding age-depth models at five locations were selected (*Supplementary Figure 2*). The selected locations are 2 km, 6 km, 8 km, 10 km, and 12 km from the shoreline. These distances corresponded to lagoonal environments, backreef, forereef, proximal slope, and distal slope. The simulated carbonate platforms accreted vertically with only minor shift of environments along the onshore-offshore axis, therefore each location can be interpreted as representing its respective environment across the entire simulation (platform lifetime). Only simulation outputs between the shore and 13 km offshore are shown in the figures, as the remainder (13-15 km) consisted entirely of redeposited carbonates and was not used further.

The simulated carbonate platforms were characterized using the following parameters calculated at any given location in the platform:

1. Stratigraphic completeness, calculated as the proportion of time steps with sediment accumulation relative to the total number of time steps in the simulation (function `get_completeness` in Hohmann et al. (2024)), corresponding to completeness on the time scale of 1 kyr (Tipper 1987; Anders, Krueger, and Sadler 1987; Kemp 2012)
2. Distribution of hiatus durations in Myr (function `get_hiatus_distribution` in in Hohmann et al. (2024)

### Simulation of phenotypic trait evolution in the time domain

We simulated three commonly discussed modes of evolution in the time domain: stasis, unbiased random walk, and biased random walk (Hunt 2006; Jones 2009; Hopkins and Lidgard 2012; Hunt, Hopkins, and Lidgard 2015). These simulations serve as “true” evolutionary history against which we compare fossil time series derived under simplified assumptions on age-depth models.

In unbiased random walk models, the change in traits over a fixed time step is independent of the previous trait values, and drawn from a distribution with mean zero (Bookstein 1987; Sheets and Mitchell 2001). In biased random walk models, the mean can deviate from zero. The trait evolution along a lineage is then generated by summing up the incremental changes in traits. Commonly used models of random walks are based on equidistant time steps, meaning the time passed between two observations of the random walk is identical for all observations. We could not use these models, because depending on where in the stratigraphic column samples are collected, the time elapsed between two successive sampling positions can vary by more than three orders of magnitude. It will be less than a thousand years when sediment accumulation rate is high, and more than half a million years when they are separated by a hiatus (*Figure 4*). If samples have identical distances in the stratigraphic column, the underlying evolutionary history will be sampled at irregular times. We use continuous-time extensions of random walk and stasis models so we can sample lineages at arbitrary time points without relying on interpolation between discrete time steps. Interpolation would potentially introduce dependencies between successive samples, contradicting the assumption of the random walk model that change in traits is independent and identically distributed. The continuous-time expansion provides exact trait values at the sampled times, and reduces to the standard discrete time step models when applied to equidistant time steps. In addition, discrete-time models incorporate time elapsed between observations into model parameters (e.g., the variance parameter of the random walk model), making their scaling behavior non-obvious when time scales are varied. Using continuous-time implementations makes it possible to compare observations across time scales and vary the granularity of observation while keeping the underlying model of evolution fixed. We use this property to examine how increased sampling of the same time interval influences model selection performance.

**Figure 4:**
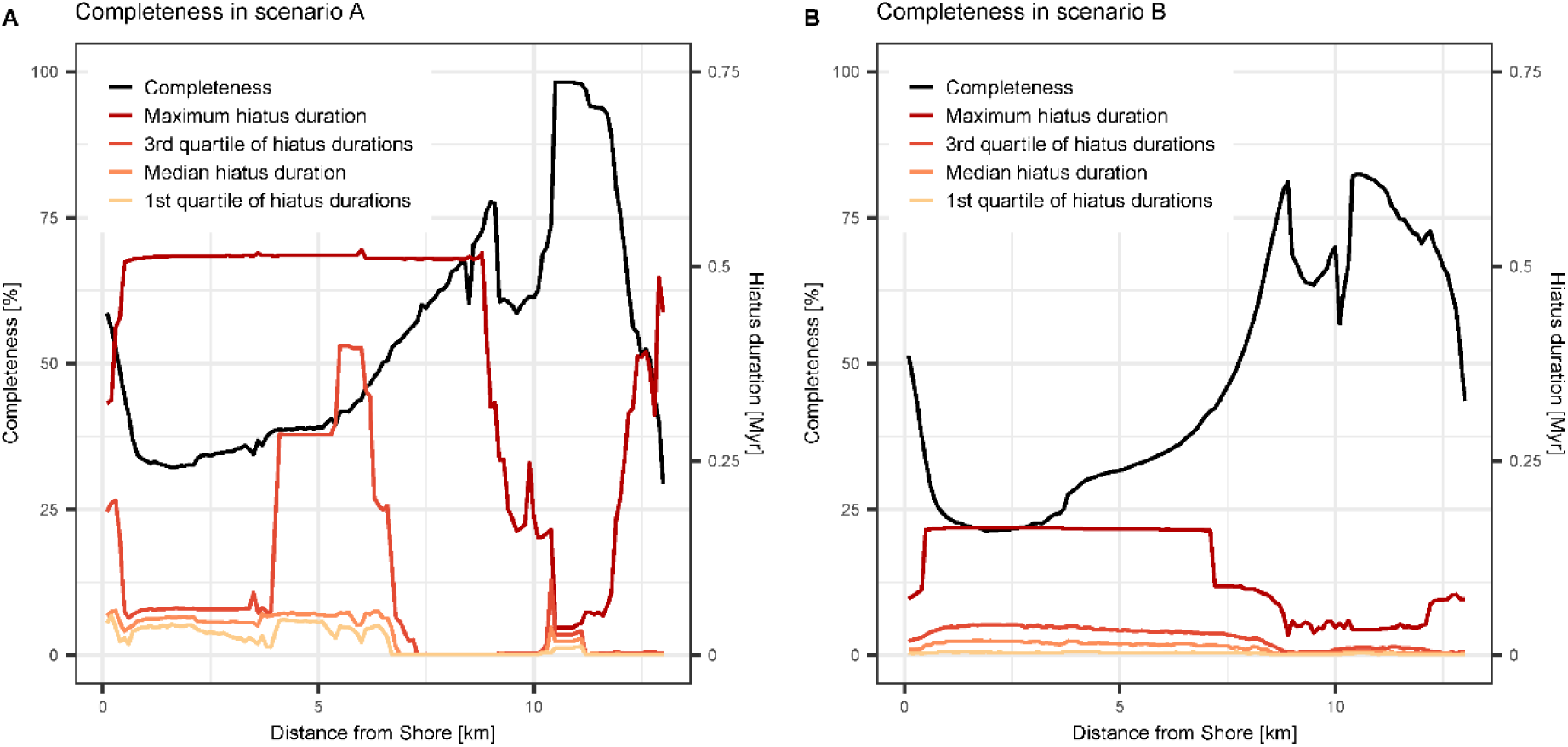
Stratigraphic completeness and distribution of hiatus durations along the onshore-offshore gradient in scenario A (left) and B (right). Maximum hiatus duration in scenario B is four times lower than in scenario B, while completeness is comparable.

We extracted age-depth models from CarboCAT Lite outputs. The age-depth models serve as functions *H : L → T* that connect the stratigraphic domain *L* with the time domain *T*. Given a model of trait evolution *M_t_* in the time domain, we then define its stratigraphic expression as *M ◦ H^-1^*, which is the trait evolution observable in the stratigraphic column. This allows us to form triplets (ℎ, *t*;, *m*), where ℎ is a stratigraphic position, *t*; is the time when that position was formed, and *m* is the trait value that describes the true evolutionary history at time *t*; and can be observed at height ℎ (*Figure 1*). The R package DAIME (Hohmann 2021b) was used for age-depth transformations to sample time and depth domain at arbitrary points. The theoretical underpinning for the need of these transformations can be found in Hohmann (2021). For the (un)biased random walk models, we use Brownian drift as a continuous-time expansion. It is given by

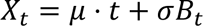

where *μ* and *σ* are model parameters, *t* is time, and *B_t_* is a Brownian motion. The parameter *μ* specifies how biased or directional the Brownian drift is and *σ* specifies how volatile the process is. After *t* time units have passed, the distribution of a trait following a Brownian drift model is normally distributed with standard deviation *σ*√*t*; and mean *μ × t + x_0_*, where *x_0_* is the trait value at the beginning of the observation. When sampled at equidistant points in time, this reduces to a standard random walk model where a normal distribution determines the step sizes.

Stasis is expressed as the lack of net change in traits over the period of observation. We model stasis as a series of independent, identically distributed random variables with mean *m* and standard deviation *s* (Sheets and Mitchell 2001). As a result, stasis is not affected by the heterodistant sampling in the time domain.

We examined the preservation of three evolutionary scenarios (*Supplementary Figure 3*):

1. Stasis with mean *m* = 0 and standard deviation *s* = 1;
2. Brownian motion as a special case of Brownian drift with parameters *μ* = 0 and *σ* = 1;
3. Brownian drift with parameters *μ* = 5 and *σ* =1, corresponding to a directional random walk.

After one million years of evolution, the Brownian motion has an expected change in traits of zero, while the Brownian drift has an expected change of traits of 5. For comparability, the *σ* parameter on the Brownian motion and drift models was kept constant, resulting in a standard deviation of traits around their mean value equal to one after one million years. The three evolutionary scenarios differ in their directionality, defined as the difference in traits accumulated over time. Brownian drift is directional, stasis and Brownian motion on average are not. Individual lineages following a Brownian motion can deviate drastically from their initial trait value – only when multiple lineages are observed, the mean trait values observed in them average to zero (*Supplementary Figure 3*).

### Sampling procedure in the stratigraphic domain

For clarity, we distinguish between stratophenetic series and time series. Stratophenetic series are taken in the stratigraphic domain and record trait values observed at specific stratigraphic positions, whereas time series are located in the time domain and record trait values observed at specific points in time.

To isolate the effects of stratigraphic incompleteness, we assume all lineages have an equal chance of being sampled across all environments. We assume identical sampling procedures for stratophenetic series in both scenarios: a sample is taken every meter and consists of 100 specimens. The distribution of traits among the specimens in a sample is normally distributed with variance of 0.1 and a mean according to the simulated trait values. This choice is made so the stratophenetic series match the format required by the paleoTS package (Hunt 2006; 2022), which we use to identify the mode of evolution. The variance of 0.1 was chosen to ensure that variability within a sample is small compared to the expected mean evolutionary change that accumulates over the course of the simulations, thus reducing the chance to mistake variability within populations for evolutionary trends (Hannisdal 2006). This procedure generates equidistant fossil time series with, depending on the location in the platform, 20 to 150 sampling positions in scenario A and 50 to 220 sampling positions in scenario B.

### Identification of the mode of evolution from simulated stratophenetic series

We applied the tests for the mode of evolution implemented in the `compareModels` function of the paleoTS package version 0.5.3 (Hunt 2022) to the simulated stratigraphic series. The tests use corrected Akaike’s Information Criterion (AICc) to determine which of the following models fits best to the stratophenetic series: the stasis model, an unbiased random walk (URW), and a biased random walk referred to as general random walk (GRW). The models identified by the `compareModels` function correspond to the simulated processes as follows: URW to the Brownian motion, GRW to the Brownian drift, and stasis to stasis. The tests take the between-samples variance in trait values and the number of specimens found at a sampling position into account. At each location in the carbonate platform, 100 stratophenetic series per evolutionary scenario (stasis, Brownian motion, Brownian drift) were simulated and tested for their fit to the stasis, URG, and GRW model. Because raw AICc values carry no meaning, we use the derived AICc weights instead. The (uncorrected) AIC weight of a model can be interpreted as the probability that it is the best approximating model, given the data and the set of candidate models (Wagenmakers and Farrell 2004). Thus, higher AIC weights (and, as a result, AICc weights) reflect better support for the model.

As a baseline for the test performance of the paleoTS package, the tests were also performed in the time domain, i.e. without any losses or distortions introduced by the stratigraphic record. To this aim, the time interval of observation (2 Myr and 2.56 Myr for scenario A and B, respectively) was subdivided into 5, 10, 15, 20, 25, 35, 50, 100 and 200 equally spaced sampling points. Lineages evolving according to the specified modes of evolution were sampled at these time points, and the test for the modes of evolution was performed on the resulting time series. Our hypothesis for this baseline is that in the absence of stratigraphic effects, increased sampling effort (i.e., a higher number of subdivisions of the time interval of observation, resulting in longer time series) increases support for the correct mode of evolution.

## Results

### Stratigraphic architectures

#### Carbonate platform A

Platform A (*Figure 2 A*; *Figure 3 A-C*) has a steep topography with high build-up up to 8 km into the basin and thin, condensed off-platform deposits between 8 and 13 km, consisting mostly of transported sediment. The majority of transported sediment is derived from the phototrophic factory, with a sharp drop in production rate at around 30 m water depth (*Supplementary Figure 1*). The thinnest interval at ca. 13 km into the basin contains also sediment transported from the two other factories.

The platform consists of two “depositional sequences” with different topographies. These “sequences” correspond to two sea-level highs, with the first one resulting in low topography and rapid progradation that led to basinward, rather than upward, growth of the platform. The second “sequence” is aggradational, with distinct “parasequences” expressed in facies, corresponding to the 125 kyr period in the sea-level curve used as simulation input. This “sequence” is responsible for the steep topography of the platform. The top of the platform is abruptly truncated as a result of the long-term cycle sea-level fall and covered with a thin deposit of the ensuing initial transgression.

The two long-term cycle sea-level falls result in two major gaps (*Figure 3 A*). The gap between the first and the second “depositional sequence” does not extend uniformly across the entire platform. During the time of the gap formation, the euphotic factory and sediment derived from transport of this factory’s products accumulated at the platform edge, prograding between 5 and 12 km basinward. The last part of this interval offlaps the platform, forming an architecture resembling the Falling Stage Systems Tract (Plint and Nummedal 2000). In contrast, the second major gap resulted in almost no deposition, reflecting almost no sediment transport in the second stage of platform formation. As a result, this platform is characterized by: two long gaps in deposition and several shorter gaps with limited spatial extent; large differences in thickness along the onshore-offshore gradient and a substantial contribution of sediment transport and offshore deposits formed entirely from sediment exported from the platform in the first half of the platform formation.

#### Carbonate platform B

For scenario B, we used the Pleistocene-Holocene global mean sea-level estimate of Miller et al. (2020). In the original dataset, the sea level varied between -121 m to 31.9 m and was here normalized to a mean of 0 m (*Figure 3 E*). The simulated time interval corresponds to gradually lowering global sea level dominated by 41-ka tilt forcing leading to amplitudes reaching 50 m. A gradual increase in sea-level amplitude throughout this interval is attributed to the onset of the glaciation of the Northern Hemisphere. In the middle Pleistocene, ca. 800 ka, a shift to “quasi-100-ka” periodicity is associated with higher sea level amplitudes reaching 140 m (Miller et al. 2020). This Milankovitch-paced sea-level changes resulted in a simulated carbonate platform with short gap durations (*Figure 3 D*, *Figure 4 B*). Where present, the gaps are widespread across the platform, i.e. sea-level drops resulted in gaps uniformly distributed across all environments. The continuity of gaps is even more pronounced in the youngest part of the platform corresponding to the last 800 kyr, reflecting the higher sea-level amplitude. Most gaps are concentrated in the central part of the platform, whereas the most shore- and basinward parts of the platform may be filled with sediment at the time when no other strata is formed in the grid, leading to the highest completeness of these two opposite ends. The dominant facies is sediment transported from the phototrophic factory (dark red in *Figure 2*), which is present across the entire platform, rather than exported off the platform as in scenario A. Hence, this platform is dominated by short-distance sediment transport. In the older part of the platform, the offshore part (at a distance of 11-13 km from the shore) has a higher stratigraphic completeness (more time represented by sediment) than the onshore end, but in the younger part of the platform, the difference disappears. The lack of large sea-level falls results in a lack of major facies shifts, low topography and a gradual, nearly aggradational buildup (*Figure 2 B*).

#### Completeness and distribution of hiatuses

Changes in completeness along the onshore-offshore gradient were very similar for both scenarios: completeness increased monotonously from the shore and had a double peak around 9 and 11 km offshore, where it reached values of 75 and 95 percent (onshore and offshore peak) in scenario A and around 80 percent (both peaks) in scenario B (*Figure 4*).

The onshore peak in completeness coincided with the forereef environment, where organisms continuously grow, while the second peak coincided with the slope onset, where transported sediment is continuously provided by the forereef. Average completeness across the platform was 53.7 % in scenario A and 46.3 % in scenario B, completeness was lowest 13 km from shore in scenario A and 1.9 km from shore in scenario B.

Qualitatively, the distribution of hiatus durations was similar for both scenarios. Maximum hiatus duration was constant over the entire platform, dropped dramatically on the slope, and then increased again off-platform. The first quartiles, medians and third quartiles of hiatus durations differed substantially from maximum hiatus durations. The exception to this is the backreef in scenario A (approx. 4–6 km from shore), where the 3^rd^ quartile of hiatus durations reaches values of 0.25 to 0.4 Myrs. Averaged over the platform, maximum hiatus duration is one to two orders of magnitude larger than median hiatus duration (135 times longer in scenario A and 13 times longer in scenario B). This shows that the distribution of hiatus durations on the platform is heavy-tailed: most hiatuses are short, but a few exceptionally long hiatuses that are associated with long-term drops in sea level are present. Quantitatively, hiatus durations in scenario A are longer (average median hiatus duration across the platform: 26 kyr in scenario A, 10 kyr in scenario B). The maximum hiatus duration is on average almost four times higher in scenario A than in scenario B.

### Stratigraphic expression of evolution

Overall, we find that stratigraphic effects on the preservation of the mode of evolution are spatially heterogeneous within the carbonate platforms (*Figure 5*), and strongly depend on (1) the directionality of the examined mode of evolution (*Figure 6)* and (2) the presence of single, long hiatuses rather than the stratigraphic completeness at the location where the stratophenetic series is sampled (*Figure 7*). As a result, under high-frequency sea-level changes (scenario B), trait evolution is much more similar to true evolutionary change in the time domain than under slow-frequency sea-level changes (scenario A).

**Figure 5:**
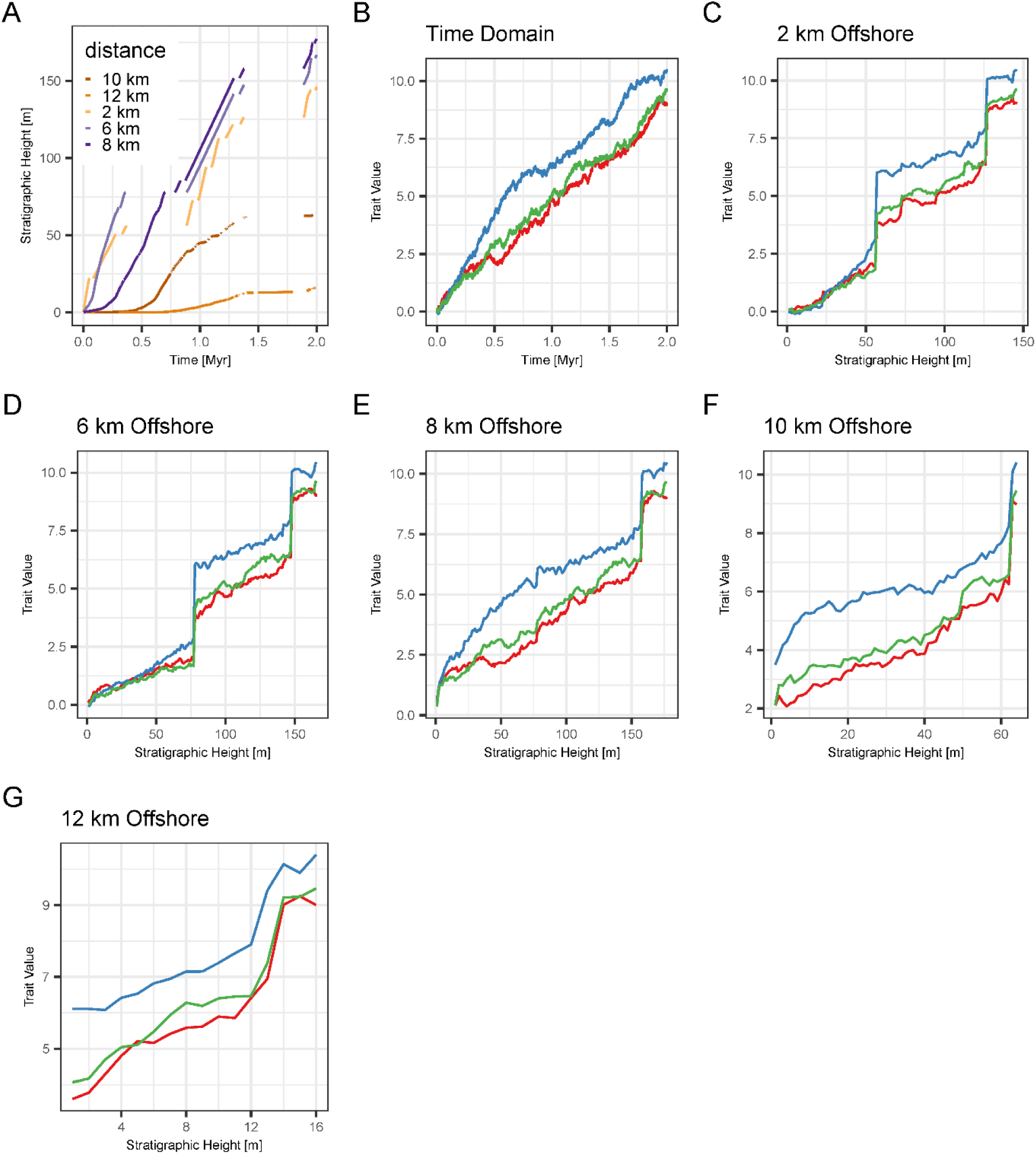
Spatial variability of the preservation of evolution in scenario A. (A) Age-depth models at varying distances from shore (B) three simulations of Brownian drift in the time domain (C), (D), (E), (F), (G) preservation of the lineages from (B) in the stratigraphic domain at 2 km, 6 km, 8 km, 10 km, and 12 km from shore in platform A. The same evolutionary history (B) is preserved differently dependent on where it is observed (C to G).

**Figure 6:**
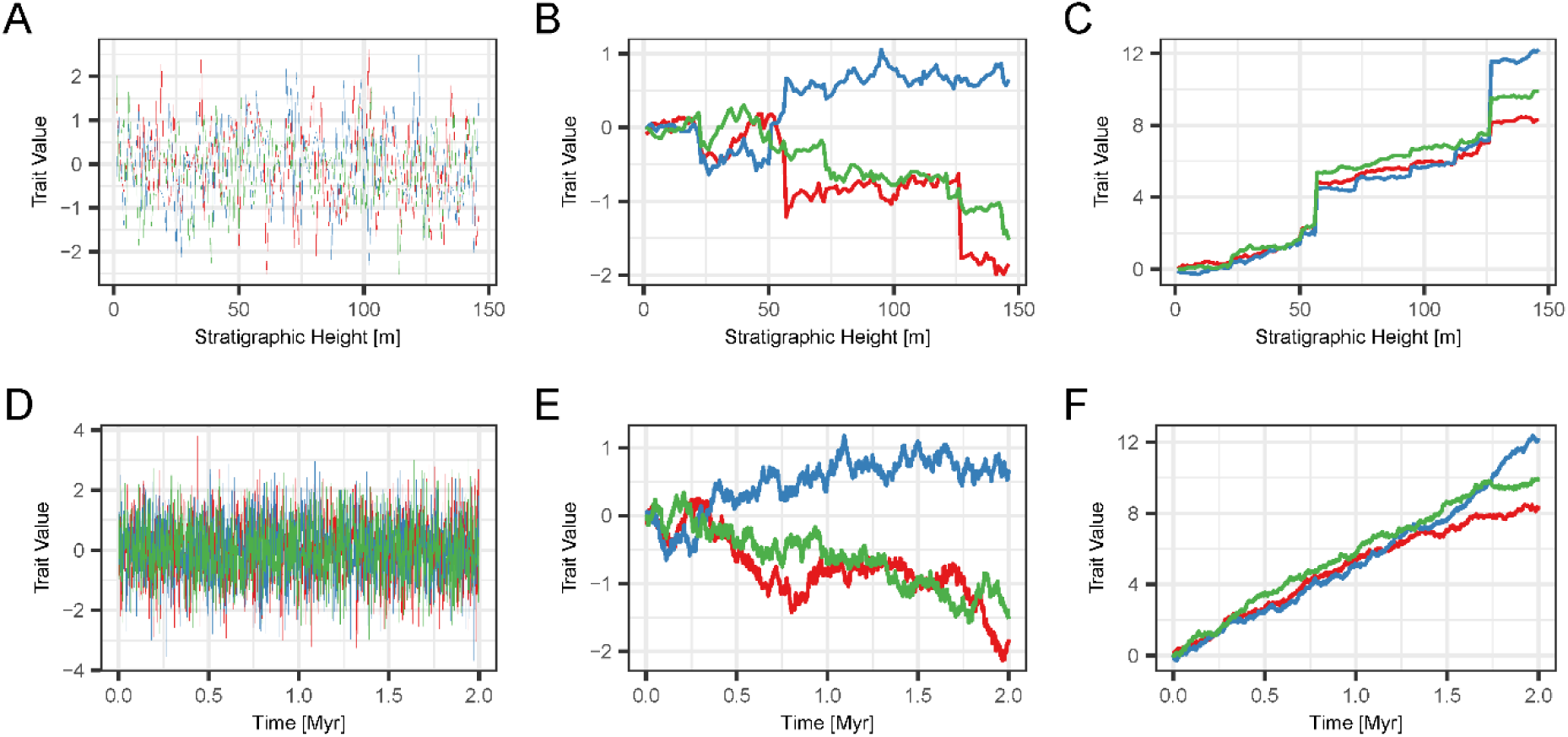
Differential preservation of different modes of evolution at the same location. First row: preservation of three lineages evolving according to the stasis (A), Brownian motion (B), and Brownian drift (C) model 6 km offshore in scenario A. Second row: The corresponding true evolutionary history in the time domain. The change in traits observable in stratophenetic series over a gap depends on the directionality of evolution and gap duration – Stasis is unaffected by the gaps, while the directional Brownian drift displays jumps in phenotype over long gaps in the stratigraphic record.

**Figure 7:**
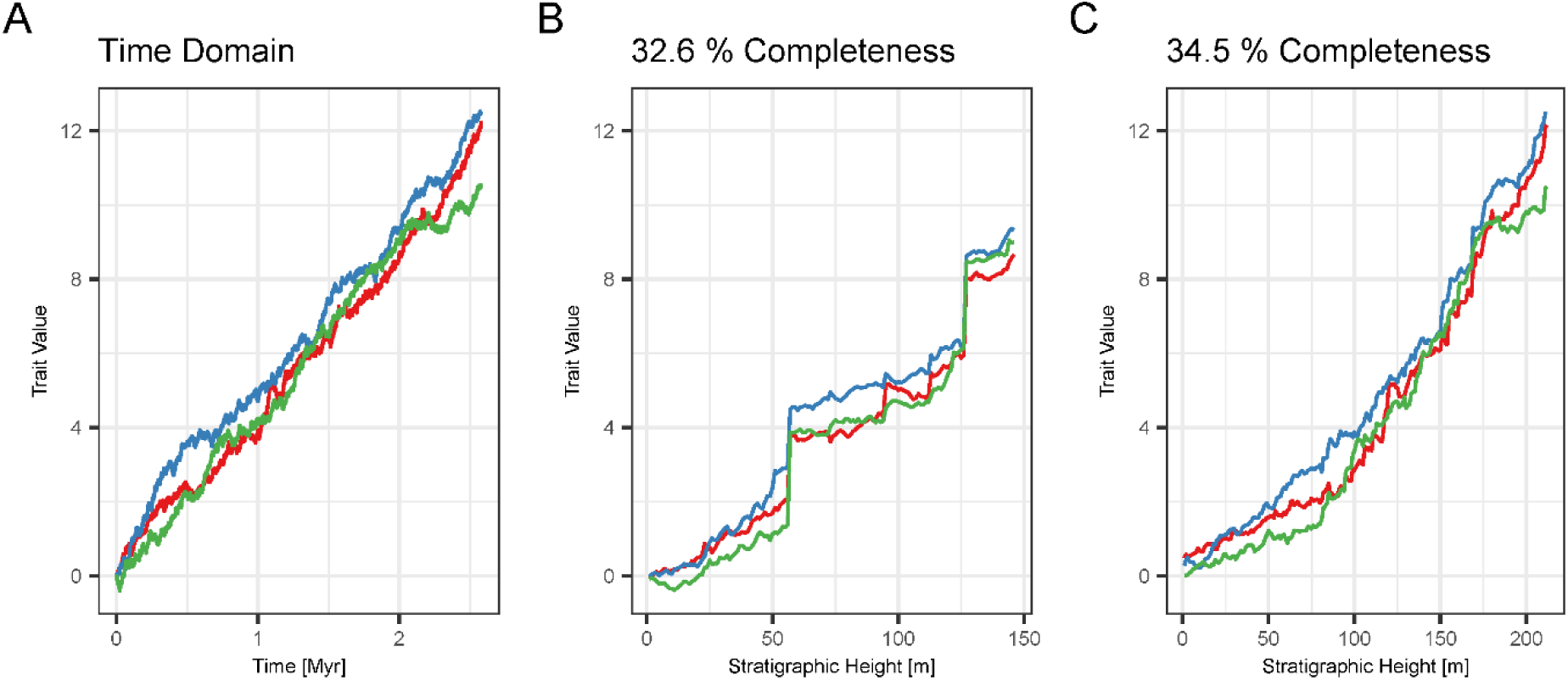
Effects of completeness vs. hiatus duration. Brownian drift in the time domain (A), 2 km offshore in scenario A (B), and 6 km offshore in scenario B (C). In these sections, stratigraphic completeness differs by only 2 %, but preservation of the lineages differs drastically due to the presence of few, but long hiatuses in scenario A generated by prolonged intervals of low sea level.

#### Differential effects of stratigraphy on modes of evolution

The extent to which stratigraphic series of trait values are affected by the architecture of the carbonate platform strongly depends on the mode of evolution (*Figure 6, Supplementary Figures 4–12*). For the same scenario and position in the platform, lineages evolving according to the stasis model are unaffected by stratigraphy. In contrast, hiatuses introduce jumps into the lineages evolving according to a Brownian drift, which are not present in the time domain. For the Brownian motion, the presence of jumps depends on whether the trait series accumulates sufficient trait differences during the hiatus, making the presence or absence of jumps over gaps effectively random (*Figure 6, B & F*, red and green vs. blue lineage). In general, jumps in traits are more pronounced when (1) evolution is more directional and (2) gaps are longer.

#### Effects of completeness

Comparing identical modes of evolution at locations in the carbonate platform with similar stratigraphic completeness shows that, perhaps counterintuitively, stratigraphic completeness is not the most important mechanism through which identification of the mode is influenced by stratigraphic architecture (*Figure 7*). If hiatuses are frequent and have similar duration, evolution in the stratigraphic domain is very similar to the time domain. If hiatuses are rare and long, traits can change significantly during the hiatus, leading to distinct differences in trait values from before to after the hiatus, generating dissimilarity between observations made in the stratigraphic domain and the “true” evolutionary history in the time domain.

#### Spatial variation

The stratigraphic expression of trait evolution varies spatially within a carbonate platform (*Figure 5*). This is a direct result of the variability of stratigraphic completeness and hiatus frequency and duration along the onshore – offshore gradient (*Figure 4*). Because of the selective effects of stratigraphic architectures on the identifiability of directional evolution, spatial variability is most pronounced for Brownian drift, and absent for stasis (*Supplementary Figures 14–18*).

### Identification of modes of evolution

Surprisingly, neither the stratigraphic architecture driven by different sea-level histories nor the location along the onshore-offshore gradient has a strong influence on the best supported mode of evolution recovered by the paleoTS package (Hunt 2022). For unbiased random walks, we found that in the absence of stratigraphic effects, for adequate models, and under excellent sampling conditions (large number of specimens, low intrapopulation variability in traits, long, equidistant time series), average support for the correct (simulated) mode of evolution decreases as time series length increases (Figure 10).

#### Test results in the stratigraphic domain

In the stratigraphic domain, the tests yielded good support for the correct mode of evolution under simulations of stasis and Brownian drift, but only intermediate support for Brownian walk. This holds for both scenarios and all locations within the carbonate platform (Figures *8 and 9*, top rows).

**Figure 8.**
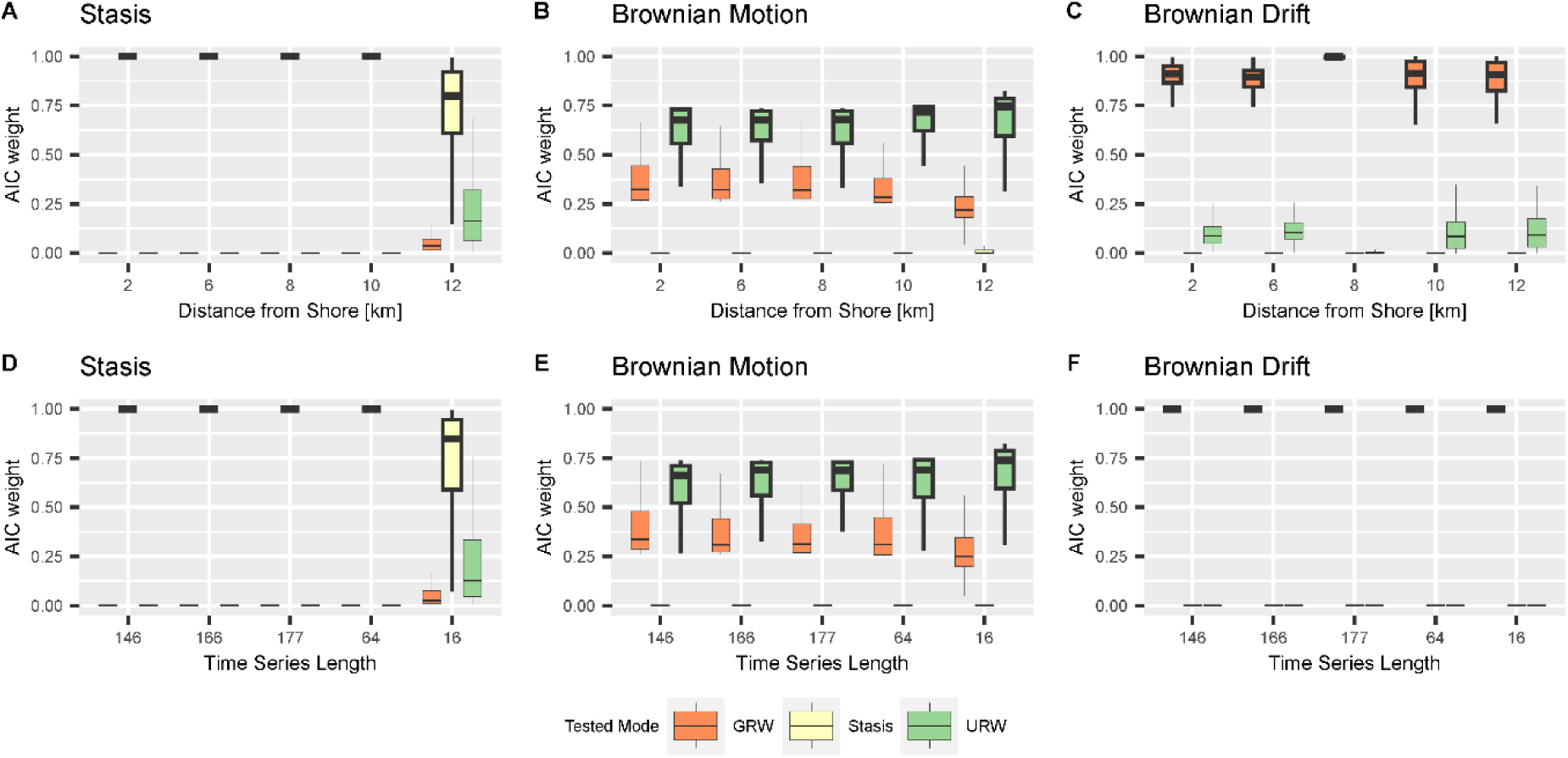
AICc weights of different modes of evolution in the stratigraphic domain in scenario A (first row, A-C) and for time series of equal length, but without stratigraphic biases (second row, D-F) under simulated stasis (first column), Brownian motion (second column), and Brownian drift (third column). The highlighted boxes are the correct test result for the simulated mode of evolution. Abbreviations for the tested modes are: GRW - general random walk, Stasis – stasis, URW – undirected random walk. Boxplot shows median and interquartile range (IQR) as hinges. Upper/lower whiskers are at most 1.5 IQR from the hinge, outliers are not shown.

**Figure 9:**
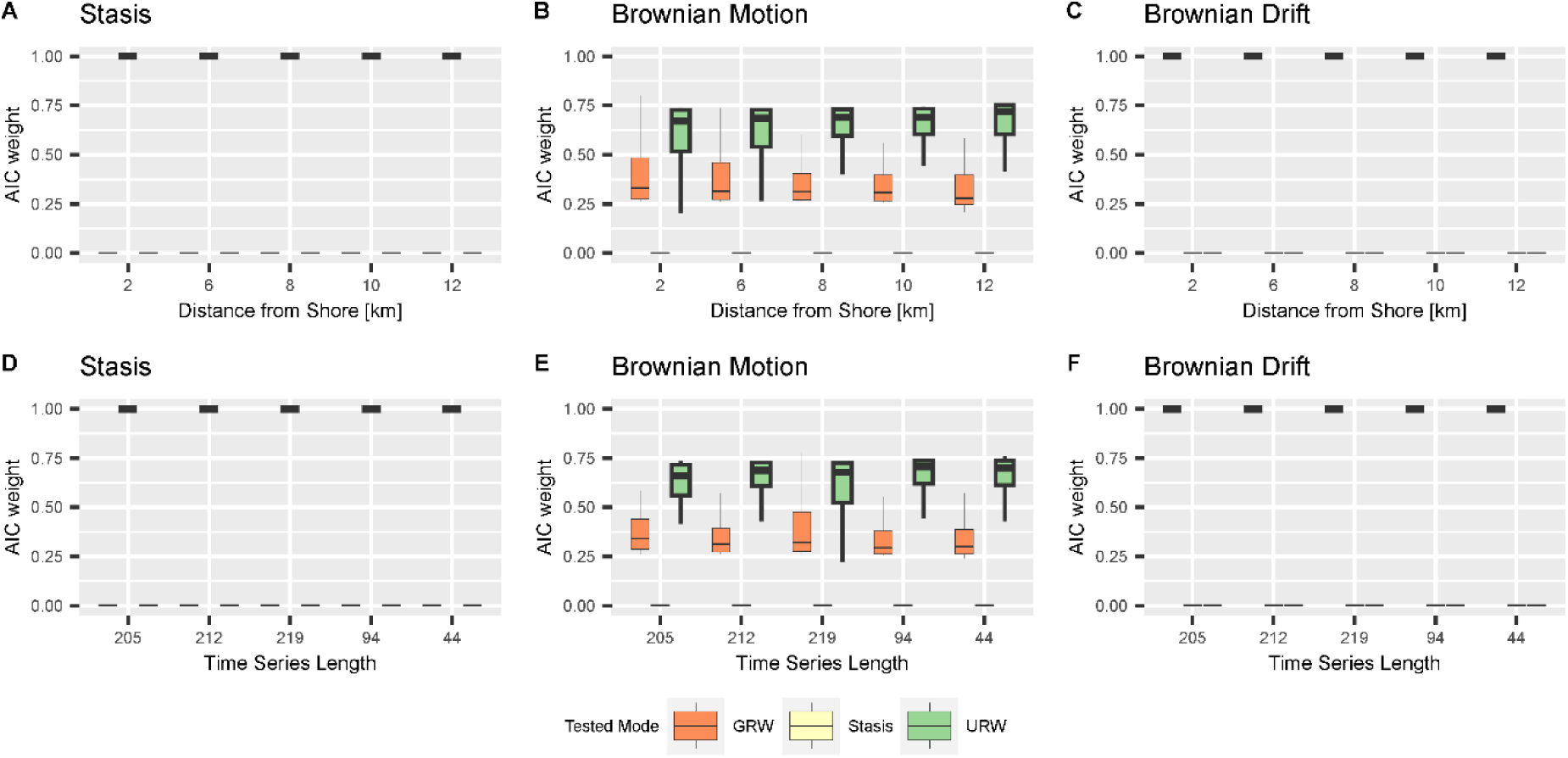
AICc weights of different modes of evolution in the stratigraphic domain in scenario B (first row, A-C) and for time series of equal length, but without stratigraphic biases (second row, D-F) under simulated stasis (first column), Brownian motion (second column), and Brownian drift (third column) at different positions in the platform. The highlighted boxes are the correct test result for the simulated mode of evolution. Abbreviations for the tested modes are: GRW - general random walk, Stasis – stasis, URW – undirected random walk. Boxplot shows median and interquartile range (IQR) as hinges. Upper/lower whiskers are at most 1.5 IQR from the hinge, outliers are not shown.

Under simulated stasis (*Figure 8 A* and *Figure 9 A*), AICc weight values for the correct mode of evolution are very high (above 0.9), except in locations where time series length is low due to a condensed record (scenario A, 12 km from the shore). Under simulated Brownian motion (*Figure 8 B* and *Figure 9 B*), AICc weights for the correct mode of evolution (URW) are moderate to high (IQR, inter-quartile range of AICc 0.6 to 0.7 in both scenarios). Support for Brownian drift is low to moderate (IQR 0.3 to 0.4 in both scenarios). This is independent of the location in the platform and of the scenario. For simulations of Brownian drift (*Figure 8 C and Figure 9 C*), average AICc weights for the correct mode of evolution (GRW) are all high across both scenarios and all locations in the platform: the IQR for scenario A is 0.87 to 0.99 and in scenario B, all AICc values are higher than 0.99. Overall, we find that neither the stratigraphic scenario nor the location in the platform (and, as consequence, stratigraphic completeness) have a strong effect on the test results.

#### Difference between the stratigraphic and time domains

After accounting for time series lengths, we find that test results with and without stratigraphic effects are very similar (*Figure 8* and *Figure 9*, comparison between the top and bottom rows). The main difference is increased dispersion of AICc weights under simulations of Brownian drift in scenario A.

#### Test performance without stratigraphic biases

Testing for the mode of evolution without stratigraphic effects (i.e., in the time domain), we found that increasing sampling resolution (time series length) from 5 to 200 sampling points per simulation time reduced support for the correct mode of evolution as measured by AICc weights under simulations of a Brownian walk, while it increased under simulations of stasis and Brownian drift (*Figure 10*, *Supplementary Figure 19*). The results for scenario A (*Figure 10*) and scenario B (*Supplementary Figure 19*) were very similar, thus we focus on scenario A here.

**Figure 10:**
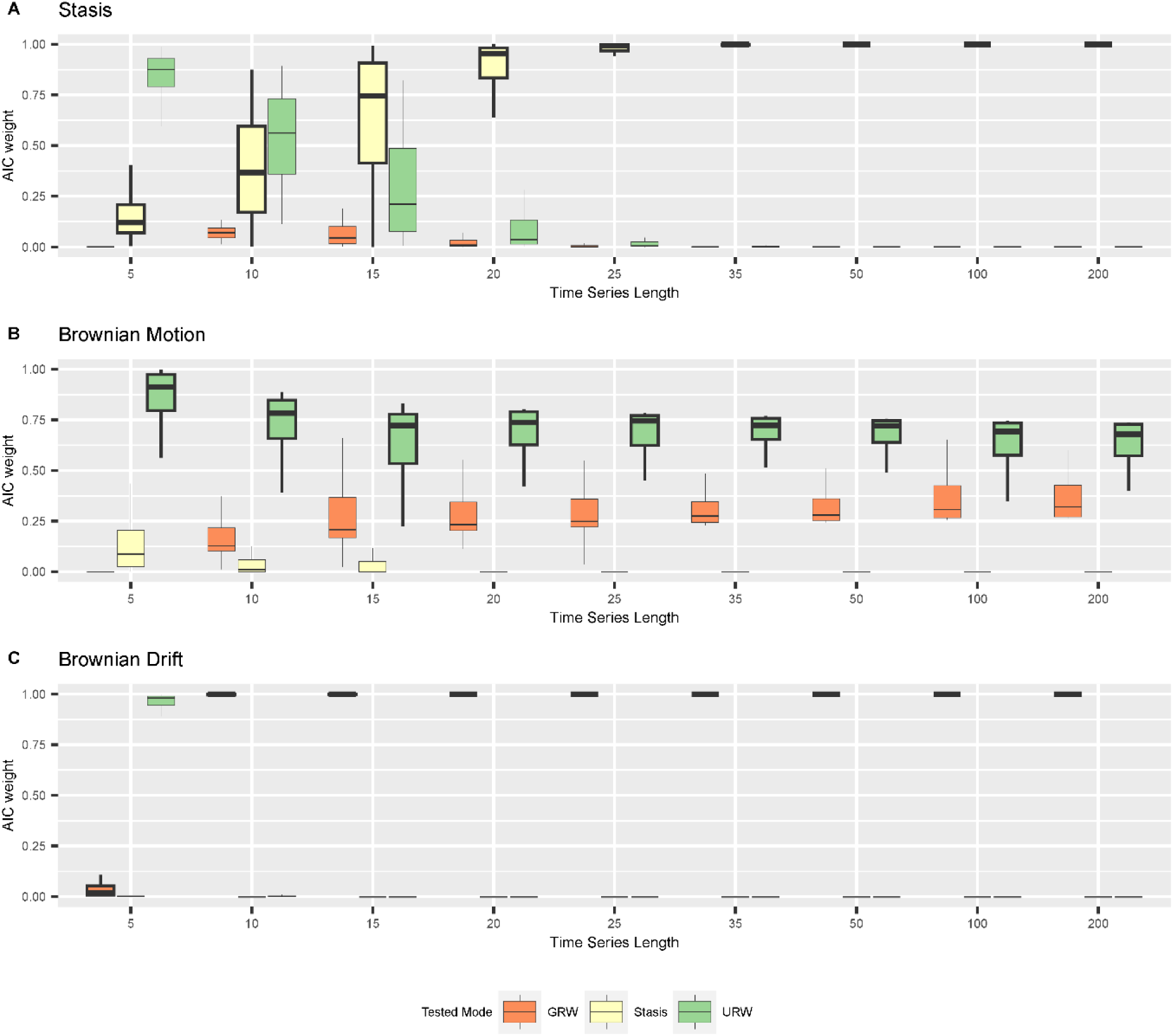
AICc weights of modes of evolution in the time domain under simulations of (A) stasis; (B) Brownian motion; (C) Brownian drift as a function of time series length. The sampled time interval is 2 Ma long (corresponding to the duration of scenario A), and is sampled with increasing frequency to reflect increasing sampling efforts. Abbreviations for the tested modes are: GRW - general random walk, Stasis – stasis, URW – undirected random walk. Boxplot shows median and interquartile range (IQR) as hinges. Upper/lower whiskers are at most 1.5 IQR from the hinge, outliers are not shown.

Under simulated stasis (*Figure 10 A*), AICc weights under low sampling effort (low number of sampling points) support URW, but this support quickly decreases and eventually vanishes for more than 25 sampling points. Support for the correct mode of evolution, stasis, is low for very short time series (5 sampling points), increases monotonously as sampling effort increases, and reaches very high support (AICc weight > 0.9) for time series with more than 25 sampling points. Support for GRW is very weak (AICc weight < 0.2) throughout. Under simulated Brownian motion (*Figure 10 B*), support for the correct mode of evolution (URW) is on average higher than for GRW, but there is substantial overlap with the range of AICc weights between both modes. For series of five samples the support for URW is the highest (IQR 0.8 to 1.0). It decreases monotonously to a mean of 0.6 (for n_samples_ = 200) as sampling effort increases. Conversely, support for both GRW increases monotonously up to a mean of 0.4 (for n_samples_ = 200) as sampling effort increases. The distributions of AICc weights for these two modes are best separated for series of 35 to 50 samples, indicating conditions under which they can be almost always distinguished. Support for stasis is low to null.

For simulations of Brownian drift (*Figure 10 C*), support for this mode is high (AICc weight > 0.9) for all time series except the very short ones (5 sampling locations), where the tests find unequivocal support for URW in all simulations.

In summary, we find that in the absence of stratigraphic effects, support for the correct mode of evolution under simulations of Brownian drift as measured by AICc weights is very good. For sufficiently long time series (more than 15 sampling points), tests correctly identify stasis, while for shorter time series, stasis can be misidentified for undirected random walk. Surprisingly, support for the correct mode of evolution (URW) decreases with sampling effort under simulations of Brownian motion (Figure 10, Supplementary Figure 19). We thus reject the hypothesis that in general, longer time series provide for better identification of the mode of evolution.

## Discussion

Simulating the preservation of trait evolution in incomplete records formed in carbonate platforms, we found that stratigraphic biases, completeness, and location in the carbonate platform have a weak effect on tests for the mode of evolution. Model support expressed by AICc weights is controlled by time series length, which is determined by the total accumulated thickness. While stasis and Brownian drift can be recovered from stratophenetic series, support for Brownian motion (undirected random walk) decreases with sampling effort. Visually, not stratigraphic completeness, but rather maximum hiatus duration and directionality of evolution determine how much series of trait values derived from the stratigraphic record differ from the true evolutionary history.

### Identifying the mode of evolution from stratophenetic series

We tested the hypothesis that the mode of evolution identified in a stratophenetic series obtained under the assumption of uninterrupted constant sediment accumulation (UCSA) is the same as the mode of the original time series in the time domain (the “true” evolutionary history). AICc weights of stratophenetic series derived under this assumption are almost identical to the AICc weights of the true evolutionary history. Stasis and directional evolution can correctly be identified from stratophenetic series under the assumption of UCSA, and AICc weights are very high (median > 0.9 for stasis, >0.67 for Brownian motion). Stasis has lower support in parts of the platform where time series are short. We attribute this to the very optimistic sampling regime we have chosen. Empirical stratophenetic series (e.g., Clyde and Gingerich (1994); Dzik (1999); Hunt and Roy (2006)) are commonly much shorter than the series simulated here. For example, stratophenetic series compiled by Hunt, Hopkins, and Lidgard (2015) had a median of 14 sampling positions and the longest stratophenetic series consisted of 114 sampling positions. In addition, we assumed a very low intrapopulation variance, and did not include any taphonomic or ecological effects that could reduce the number of specimens. Hannisdal (2006) argued that these effects combined can lead to “analytical stasis”, where the mode of evolution cannot be distinguished from stasis due to low statistical power, potentially explaining the abundance of morphological stasis observed in the fossil record (Hunt, Hopkins, and Lidgard 2015). Due to our sampling strategy, we could not confirm this effect, but found an opposite effect in the absence of stratigraphic biases: under simulations of stasis, tests favor the recognition of unbiased random walks for very short time series (less than 10 sampling points) and provide mixed support for both stasis and unbiased random walks for short time series (15 sampling points) (*Figure 10A).* For simulations of Brownian motion, tests find intermediate to good support for the correct mode of evolution (URW, unbiased random walk), which decreases with time series length even in absence of stratigraphic biases (Figures *8* and *9*, lower rows, *Figure 10*). This effect is not due to our implementation of trait evolution, as it persists when the internal simulation procedures provided by the paleoTS package are used (Hohmann and Hopkins 2024). Based on the probabilistic interpretation of the AICc weights, this implies that the more samples are collected, the lower the probability that the data-generating model is the best model (Wagenmakers and Farrell 2004). There are multiple potential explanations for this apparent statistical paradox. One is that sampling a fixed time interval more frequently leads to diminishing returns on the information gain per sample. Intuitively, a few samples should be sufficient to separate directional evolution from stasis. However, this does not explain why the effect only appears under simulations of Brownian motion, and AICc weights for the correct mode of evolution decrease instead of increasing. Another potential explanation is that AIC is not consistent in the sense that if sample size grows, the probability that AIC identifies the true model does not approach one (Bozdogan 1987). It is unclear if the considered time series are long enough to display such asymptotic effects, and how they translate from AIC to AICc weights. The decrease in AICc weight might be caused by the fact that the models tested for are nested in the sense that URW is a special case of GRW, making it impossible to strictly separate between them. Ornstein-Uhlenbeck (OU) processes are used to model trait evolution towards an optimum. They have multiple modes of evolution nested within them: with no selection, they reduce to an unbiased random walk; when they start in the phenotypic optimum, they represent (autocorrelated) stasis; and when the optimum is far away they represent directional evolution (Cooper et al. 2016; Grabowski et al. 2023). If OU is included in the set of tested modes of evolution, support for them increases with sampling effort, independent of the mode of evolution simulated (Hohmann and Hopkins 2024). This is an indicator that the decrease in AICc weights for URW under simulations of Brownian motion is driven by nestedness of models. This highlights that the separation into distinct modes of evolution is conceptually useful, but mathematically imprecise and needs to be approached with caution.

Lastly, individual sample paths of a Brownian walk can look directional (*Figure 6 E)* as a result of random fluctuations. All models of unbiased random walks have an expected trait value of 0. However, the probability of deviating from 0 increases with time, potentially creating lineages with large deviations from the expected trait value. For the unbiased random walk model, the evidence ratio (ratio of AICc weights (Portet 2020)) of individual simulations can be close to 1, making unbiased and biased random walks equally fitting models. In our simulation study, we have access to the distribution of AICc weights under repeated sampling of the underlying mode of evolution. Supplementary Figure 20 illustrates the distribution of the evidence ratio (the ratio of AICc value for the correct mode, URW, to the AICc value for GRW) for one of the cases where the distributions of AICc weights overlap substantially, i.e. for the set of simulated Brownian motion in scenario A at a distance of 6 km from the shore. In 18% of runs, the incorrect (GRW) mode has higher support than the correct one, and the AICc for the wrong mode can be as high as 0.959. Overlaps between the AICc distributions of alternative models are also observed for very short time series, e.g. between stasis and URW (*Figure 10 A*), pointing to situations where incorrect decisions on single trait sequences are more likely.

The simulation approach opens the possibility to estimate how common a wrong decision would be made as a result of the variable properties of any single run. However, for empirical studies, only one lineage representing one sample path or realization of the underlying evolutionary process is known. Based on the resulting point estimate of the AICc weight, unbiased and biased random walk might not be distinguishable. In the entire study, Brownian motion was the mode most susceptible to be mis-identified as Brownian drift, which coincides with the (not always correct) intuition that strongly directional series must be adaptative. Our results provide a guideline on when it is the case and an empirical estimation of the frequency of false decisions when single trajectories are investigated.

### Effects of stratigraphic incompleteness

The second hypothesis we examined was that the lower the stratigraphic completeness, the lower the chance to identify the correct mode of evolution from stratophenetic series that are constructed based on the assumption of uninterrupted constant sediment accumulation (UCSA). Spatial variability in AICc weights in both carbonate platforms can be explained by differences in time series length resulting from differences in total accumulated thickness (compare Figures *8* and *9* with *Figure 10*) rather than differences in stratigraphic architectures or completeness. One possible explanation for the high similarity of test results in the time and stratigraphic domain is that they are based on the corrected Akaike Information Criterion (AICc), which measures relative fit of models to data. In the time domain, we simulate lineages according to the same model we test for. For Brownian walk and drift, the modes of evolution tested for are not adequate to describe the trait evolution in the stratigraphic domain as they do not incorporate any jumps. For example, the stratigraphic expression of the Brownian drift models resembles a Lévy process with rare, but large, jumps, rather than a Brownian drift (Landis, Schraiber, and Liang 2013; Landis and Schraiber 2017) (*Figure 5*). The issues arising from using AICc when models are not adequate are well known, and the adequacy of models of phyletic evolution has been discussed by Voje, Starrfelt and Liow (2018) and Voje (2018). Despite the lack of model adequacy in the stratigraphic domain when UCSA is assumed, paleoTS test results are not affected by the non-linear transformation of time series by age-depth models.

In addition to the tests, we visually compared stratophenetic series derived under this assumption with time series of the true evolutionary history (*Figures 5, 6, 7, Supplementary Figures 4—19*). Trait evolution reconstructed using simplified age-depth models varies spatially throughout the platform (*Figure 5*). Even if gaps are identified and their duration is known, evolutionary history coinciding with sea level drops is not preserved on the platform top. These time intervals can be recovered from the distal slope where sediment is continuously accreted during lowstand shedding (Figures *3* and *4*)(Grammer and Ginsburg 1992). This highlights the importance of combining information from across the entire sedimentary basin in reconstructing past changes (Holland and Patzkowsky 2015; Zimmt et al. 2021), although the information gained from this approach might vary between carbonate platforms and siliciclastic systems.

We found that stratigraphic incompleteness (Dingus and Sadler 1982; Tipper 1987) is an imperfect predictor of the biasing effects of stratigraphic architectures on phenotypic evolution. For similar values of stratigraphic completeness, the stratigraphic expression of the same lineage can strongly differ. For example, stratophenetic series of Brownian drift reflect the underlying true evolutionary history, or display apparent jumps in phenotype for very close values of completeness (*Figure 7*). In addition, different modes of evolution are biased to a different degree when compared within the same section (and thus identical stratigraphic effects and values of incompleteness). For example, in the same section, Brownian drift displays jumps in phenotype over gaps that will be misinterpreted as elevated rates of evolution, whereas stasis remains unaffected (*Figure 6*). Multiple definitions of incompleteness have been applied in stratigraphy (Anders, Krueger, and Sadler 1987), some of which incorporate spatial variability (Straub and Foreman 2018). The definition of stratigraphic completeness from Tipper (1987) we used here reflects the intuition that gaps in the geological record are the dominant factor that diminishes the quality of the fossil record. While other definitions of incompleteness might be more suitable to quantifying stratigraphic effects, our results show that the nature of the underlying evolutionary process cannot be neglected when assessing the fidelity of the fossil record.

The change of phenotype over a gap is determined by gap duration and the change in phenotype accumulated over this time interval. Stasis remains unaffected by gaps as it accumulates no change in phenotype. On the other hand, for Brownian drift, change in phenotype is proportional to gap duration. Under the simplified assumptions on ADMs we used, the change in traits over gaps will be taken at face value, giving trait evolution a distinct punctuated look (Rita et al. 2019). In paleontological, and especially biostratigraphic, practice, a sudden change in a trait is typically the basis for recognizing a new species and, thus, a speciation event. It would not meet the original definition of punctuated equilibrium (Gould and Eldredge 1972), which requires the speciation to take place allopatrically.

However, a precise modern formulation of the punctuated equilibrium hypothesis is debated (see Pennell, Harmon, and Uyeda (2014) vs. Lieberman and Eldredge (2014)), and identifying allopatry and the type of speciation relies on geological and morphological information. This creates the risk of circular reasoning in identifying whether a given sudden change should be considered a case of punctuated change (e.g. sympatric speciation) or punctuated equilibrium. Models of trait evolution that can incorporate punctuations are Lévy processes, a class of stochastic process that combines gradual change with discrete jumps (Landis, Schraiber, and Liang 2013; Landis and Schraiber 2017). In Lévy processes, the jump components are random and have a Poisson structure. This provides a way to separate between artefactual jumps introduced by simplistic ADMs and true elevated rates of evolution: Artefactual jumps will coincide with gaps in the record or interval of reduced sediment accumulation rates, and thus connect to external drivers of stratigraphic architectures such as drops in sea level, rather than being random. Our study provides the stratigraphic null hypothesis that punctuated change in morphology should be more prevalent at times with low frequency sea-level changes. Combined, our results suggest that due to gaps in the stratigraphic record, stratophenetic series favor the recognition of both stasis and complex, punctuated models of evolution (Hunt, Hopkins, and Lidgard 2015).

Hoffman (1989) pointed out that most paleontological evidence for punctuated equilibrium comes from shallow water habitats (e.g. (Kelley 1983; Williamson 1981)), which he argues are more incomplete and thus favor the recognition of punctuations. This reflects the common idea that different environments have different incompleteness and thus different abilities to preserve evolution, as we have phrased in our hypothesis two. In contrast to this preconception, we found that many environments in a carbonate platform have very similar incompleteness and hiatus distributions (*Figure 4*). Due to the flat geometry of the platform, the separation is between platform top and slope, not the distinct environments within the platform such as forereef and lagoonal environments (Liu and Liu 2021). In addition, incompleteness alone is not sufficient to produce artefactual jumps in phenotype (*Figure 7*).

Gaps in the stratigraphic record need to be sufficiently long compared to the rate of the evolutionary process, so that morphologies can change enough to display sufficient offset over the gap. This indicates that, rather than incompleteness, maximum hiatus duration is driving the discrepancy between stratophenetic series and the true evolutionary history.

Naturally, maximum hiatus duration is limited by stratigraphic completeness and the time covered by the section. But even under high incompleteness, as long as hiatus frequency is high and durations are short, the fossil record can still give a good insight into evolutionary history, and be adequate to test a wide range of evolutionary hypotheses (Paul 1992).

In our stratigraphic models, hiatus frequency and duration are determined by the frequency of sea-level fluctuations. Preservation of evolutionary history under high-frequency sea level changes (scenario B) is good, while the large-scale fluctuations in scenario A result in prolonged gaps with a large impact on the recovery of the mode of evolution (Figure 7). This demonstrates that understanding controls on the spatial and temporal heterogeneity of the stratigraphic record is decisive in correctly interpreting the evolutionary history of a lineage in the realistic case of spatially constrained sampling.

### Limitations of the simulated carbonate platforms

Carbonate platforms used for this research only approximate stratigraphic architectures that would form in nature. First, CarboCAT Lite does not include erosion other than sediment transport, which removes sediment locally immediately after its formation, but preserves the total volume of generated sediment in the platform. In real carbonate platforms, part of the carbonate sediment is dissolved and returned to the water column (e.g., Albright, Langdon, and Anthony (2013), exported into the ocean, and a part is chemically and mechanically eroded when it becomes emerged. The addition of erosion to the model would have the effect of enlarging the present gaps in the record and merging the shorter ones, resulting in fewer, longer gaps and lower stratigraphic completeness. Second, carbonate production curves used to inform the models reflect activities of carbonate factories under stable conditions. In reality, a regression would often lead to removal of carbonate producing organisms from the emerged area. The recolonization of this area by carbonate producers would lead to a lag in the factory resuming its activity. Carbonate production and preservation is also sensitive to diurnal, seasonal and long-term astronomical cycles, e.g. through temperature control over the proportion of precipitated aragonite (Balthasar and Cusack 2015) and the length of the vegetative season of calcifiers (e.g., Marshall and Clode 2004; Mancuso et al. 2019).

Our simulations did not include pelagic carbonate production, spatially heterogeneous subsidence or gravitational sediment transport, which may be crucial in the formation of some empirical carbonate platforms (Masiero et al. 2020). We relied here on high benthic *in situ* production rates as the dominant driver of sediment accumulation in tropical, attached carbonate platforms. Thus, the stratigraphic architectures presented here are conceptual endpoints with exaggerated completeness.

The construction of our study follows the assumption of uninterrupted constant sediment accumulation (UCSA), which implies that the ages of fossils found at a given stratigraphic position correspond to the age of the stratum. This assumption is made in many simulation studies of trait evolution in the fossil record (e.g. (Hannisdal 2006)) and inherent to most methods that estimate age-depth models (e.g. (Parnell et al. 2008)). With this respect, our computer experiment is representative for such studies. However, time averaging and sedimentary condensation mean that typically more time is represented by fossils than by sedimentary strata (Kowalewski and Bambach 2008; Tomašových et al. 2022). On the other hand, fossils of different ages will be found at the same stratigraphic height, limiting the temporal resolution of evolutionary steps in the population’s mean. Because of high *in situ* sediment production, timescales of time averaging identified in modern tropical carbonate environments are typically much shorter than rates of trait evolution, regardless of the way they are measured (Hunt 2012; Voje 2016; Kowalewski et al. 2017; Philip D. Gingerich 2019). This means that trait evolution reconstructed in tropical carbonate platforms may not be representative for other environments with lower rates of sediment accumulation or higher proportion of transported material.

For simplicity, we assumed that simulated taxa are equally represented and preserved across all sampled environments. Specifying their environmental niche or differential preservation would further reduce the number of data points in stratophenetic series. Holland (2000) showed that the record of a stenotopic species would be even more incomplete than the simulations presented here.

## Conclusions

We tested the hypothesis that the commonly employed approach to identifying the mode of evolution in fossil succession, i.e. linear projection of stratigraphic positions of occurrences into the time domain without considering changes in sedimentation rate and gaps in the record, recovers the correct mode of evolution. We found that, although prolonged gaps distorted trait evolution record visually, tests for the mode of evolution were only weakly affected by gaps and irregular age-depth models. Our findings differ from those of Hannisdal (2006), who found (using a different approach but asking the same question) that incomplete sampling in the stratigraphic record may result in all other modes of evolution being identified as stasis.

Our findings did not vary substantially between two stratigraphic architectures with varying gap distributions and degrees of stratigraphic completeness. The lack of influence of stratigraphic effects is counterintuitive, as deeper environments are often assumed to be more complete and therefore more suitable for sampling fossil series for evolutionary studies. In the case of undirected random walks, increasing the number of observations (i.e. sampling intensity, length of the fossil series) did not improve the identification of the mode of evolution, but rather worsened it.

Our study was motivated by improving the recovery of evolutionary information from highly resolved fossil successions, particularly at microevolutionary scales. We are convinced that such successions can aliment models and understanding that is not accessible to exclusively neontological methodologies, as illustrated by e.g. Hopkins and Lidgard (2012); Voje (2016); Petryshen et al. (2020). Our contribution is the use of stratigraphic forward modeling to ground-truth the methodologies serving this palaeobiological research program. Forward modeling allows rigorous testing of concerns that the fossil record is too distorted, or too incomplete, to answer (micro)evolutionary questions. They also offer the possibility to evaluate the robustness of identifications of the mode of evolution and estimates of its parameters under different simulated age-depth models. Finally, sedimentological information may aliment age-depth models (Hohmann 2021a; Jarochowska et al. 2020). The proliferation of ever better stratigraphic forward models (e.g. CarboCAT (P. M. Burgess 2013; Masiero et al. 2020), SedFlux (Hutton and Syvitski 2008), strataR (Holland 2022), CarboKitten.jl (Hidding et al., 2024)) opens the possibility to validate these methods and improve our understanding of the fossil record.

## Acknowledgements

The authors would like to thank Melanie Hopkins, Katharine Loughney, Bjarte Hannisdal, Kenneth De Baets, Gene Hunt, and one anonymous reviewer for their feedback on the manuscript. The authors would also like to thank the Peer Community In (PCI) Paleontology.

## Declarations

### Ethics approval and consent to participate

Not applicable

### Consent for publication

Not applicable

## Availability of data and materials

The dataset supporting the conclusions of this article is available in the Zenodo repository, https://doi.org/10.17605/OSF.IO/ZBPWA (Hohmann et al. 2023). All code used can be found in (Hohmann et al. 2024) and is accessible via https://doi.org/10.5281/zenodo.11394049 . Supplementary figures can be found in the supplementary materials.

### Competing interests

The authors declare that they have no competing interests

## Funding

Funded by the European Union (ERC, MindTheGap, StG project no 101041077). Views and opinions expressed are however those of the author(s) only and do not necessarily reflect those of the European Union or the European Research Council. Neither the European Union nor the granting authority can be held responsible for them.

## Authors’ contributions

According to the CRediT taxonomy

**Niklas Hohmann:** Conceptualization, Methodology, Software, Validation, Formal analysis, Investigation, Data curation, Writing – Original Draft, Writing – Review & Editing, Visualization.

**Joël R. Koelewijn:** Software, Validation, Formal analysis, Investigation, Visualization.

**Peter Burgess**: Writing – Review & Editing, Software.

**Emilia Jarochowska:** Conceptualization, Methodology, Validation, Visualization, Writing – Review & Editing, Supervision, Project administration, Funding acquisition.

## Notes

https://doi.org/10.17605/OSF.IO/ZBPWA

https://doi.org/10.5281/zenodo.11394049

### Competing Interest Statement

The authors have declared no competing interest.

### Summary of Updates

Changed layout after acceptance for recommendation in PCI paleontology

