## supplementary figures for "Identification of the mode of evolution in incomplete carbonate successions" (from another version of this preprint: version 3, 2024-05-30)

### Identification of the Mode of Evolution in Incomplete Carbonate Successions – Supplementary Figures

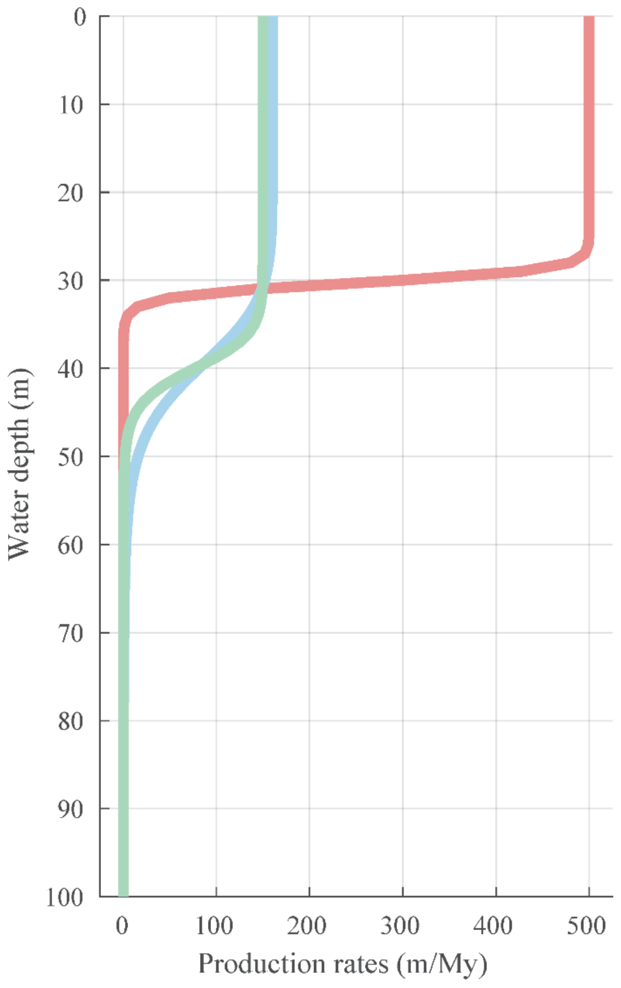


*Supplementary Figure 1: Carbonate production for the three producing facies as a function of depth.*

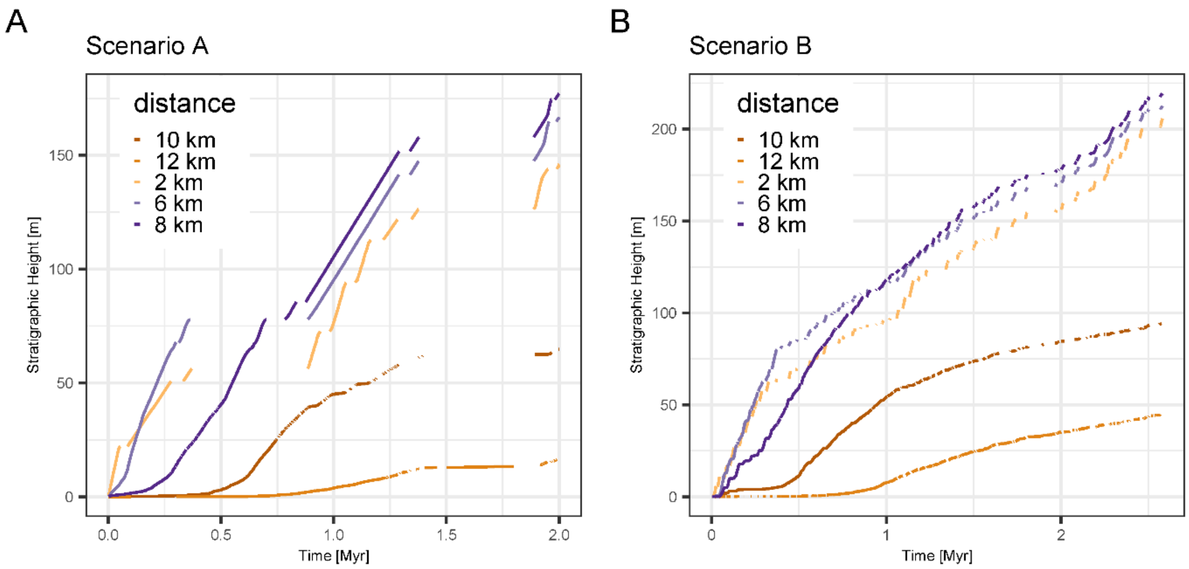


*Supplementary Figure 2: Age-depth models at the examined locations in scenario A and B.*

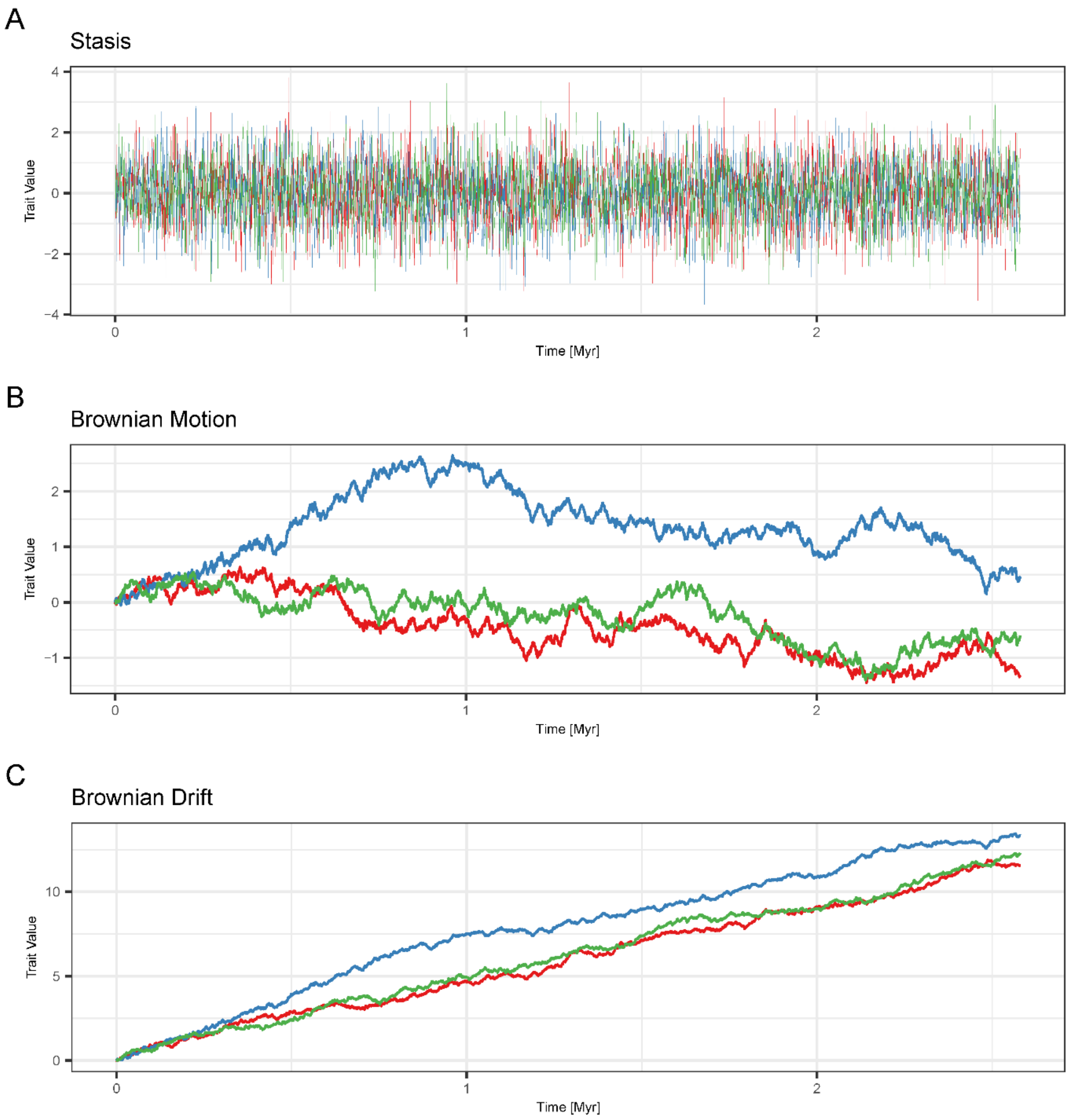


*Supplementary Figure 3: Simulations of the three examined evolutionary scenarios in the time domain.*

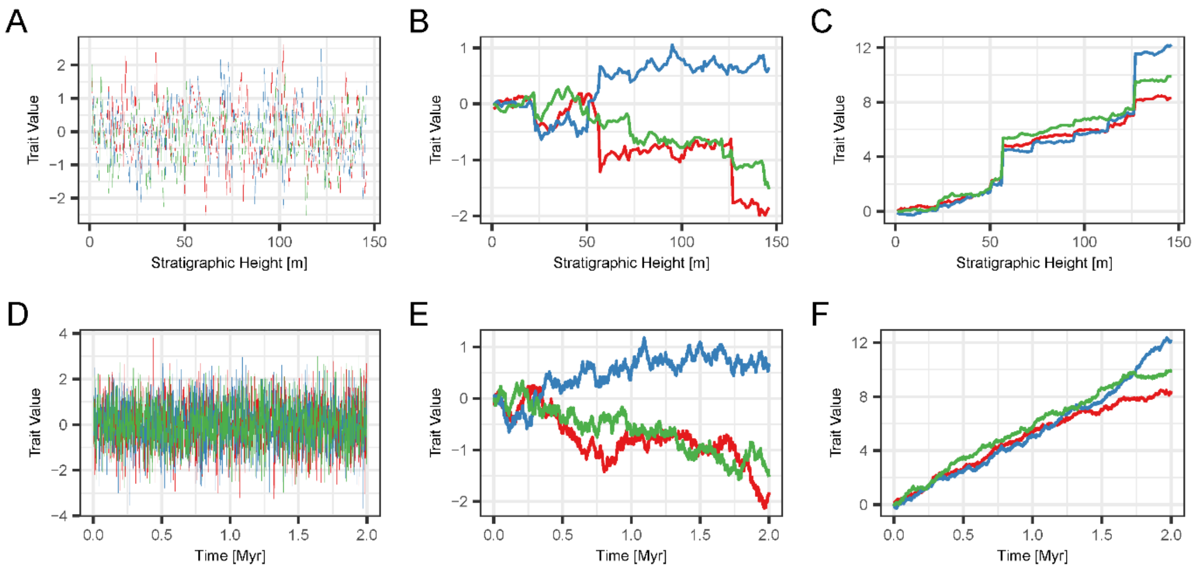


*Supplementary Figure 4: Differential preservation of different modes of evolution in scenario A, 2 km from shore. First row: preservation of three lineages evolving according to the stasis (A), Brownian motion (B), and Brownian drift (C) model. Second row: The corresponding true evolutionary history in the time domain.*

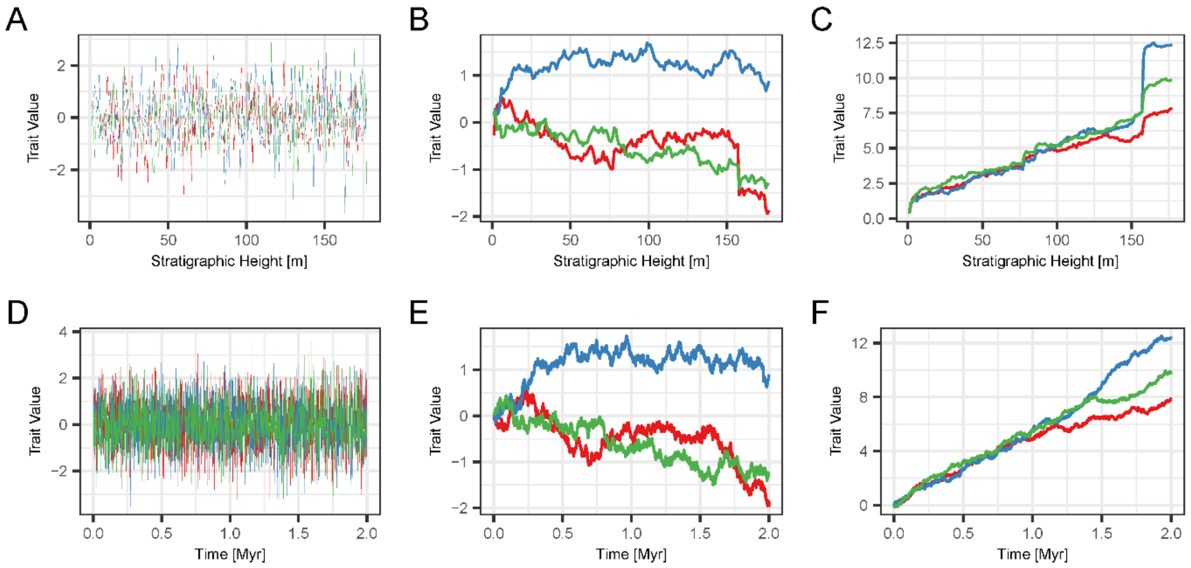


*Supplementary Figure 5 Differential preservation of different modes of evolution in scenario A, 8 km from shore. First row: preservation of three lineages evolving according to the stasis*

*(A), Brownian motion (B), and Brownian drift (C) model. Second row: The corresponding true evolutionary history in the time domain.*

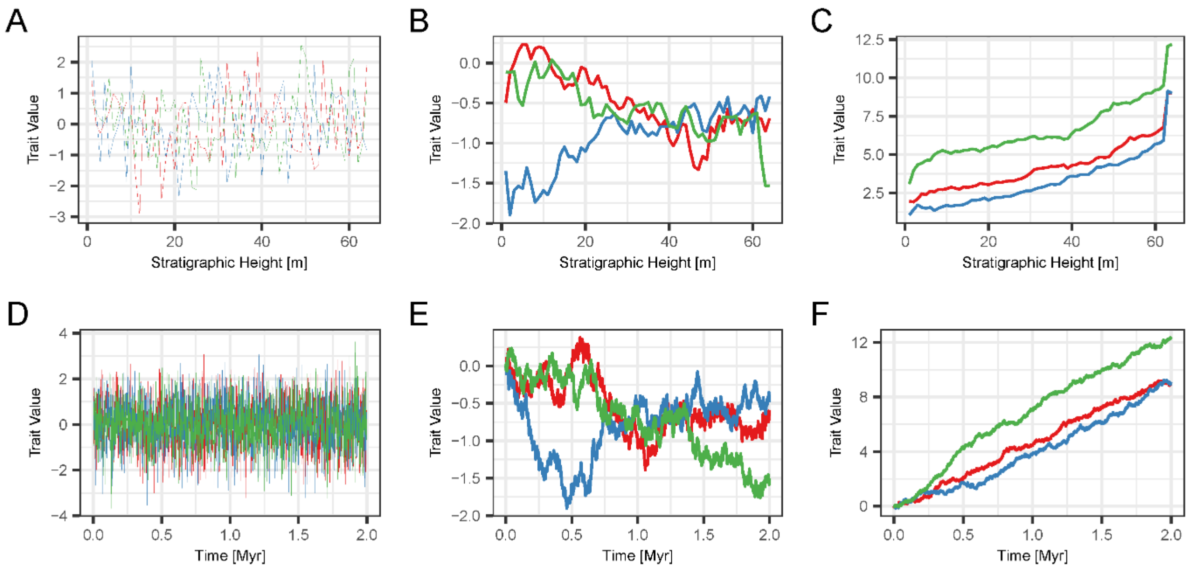


*Supplementary Figure 6: Differential preservation of different modes of evolution in scenario A, 10 km from shore. First row: preservation of three lineages evolving according to the stasis (A), Brownian motion (B), and Brownian drift (C) model. Second row: The corresponding true evolutionary history in the time domain.*

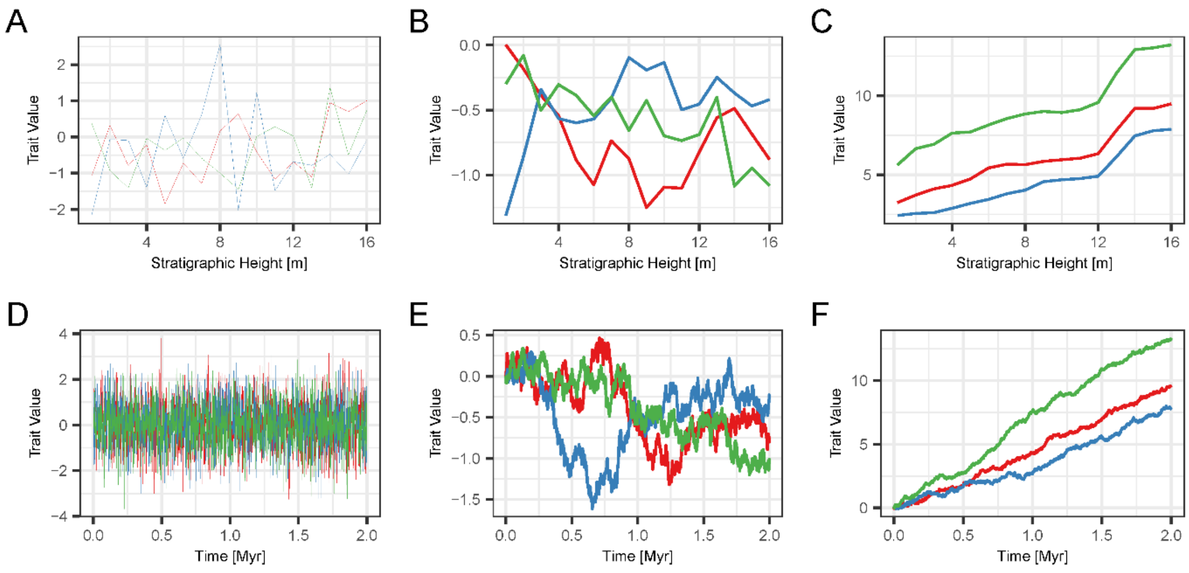


*Supplementary Figure 7 Differential preservation of different modes of evolution in scenario A, 12 km from shore. First row: preservation of three lineages evolving according to the stasis (A), Brownian motion (B), and Brownian drift (C) model. Second row: The corresponding true evolutionary history in the time domain.*

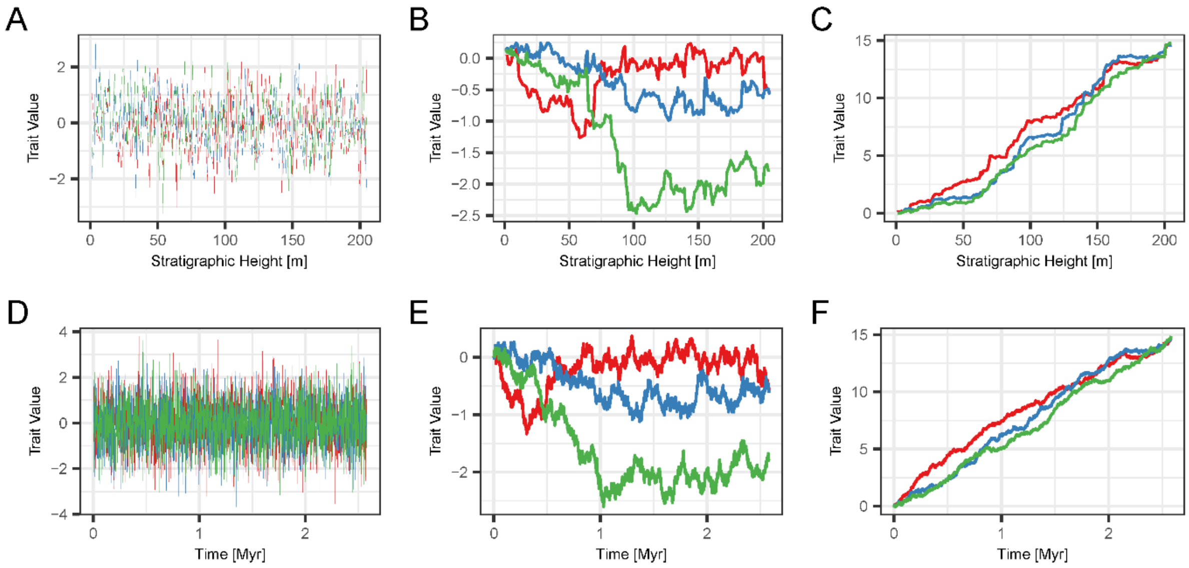


*Supplementary Figure 8: Differential preservation of different modes of evolution in scenario B, 2 km from shore. First row: preservation of three lineages evolving according to the stasis*

*(A), Brownian motion (B), and Brownian drift (C) model. Second row: The corresponding true evolutionary history in the time domain.*

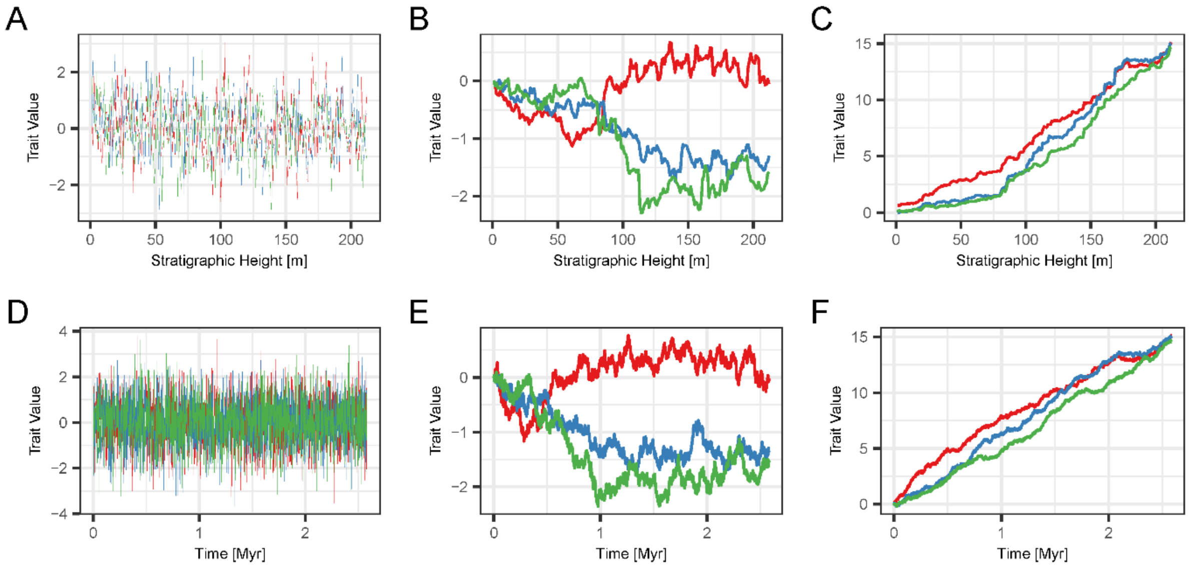


*Supplementary Figure 9: Differential preservation of different modes of evolution in scenario B, 6 km from shore. First row: preservation of three lineages evolving according to the stasis (A), Brownian motion (B), and Brownian drift (C) model. Second row: The corresponding true evolutionary history in the time domain.*

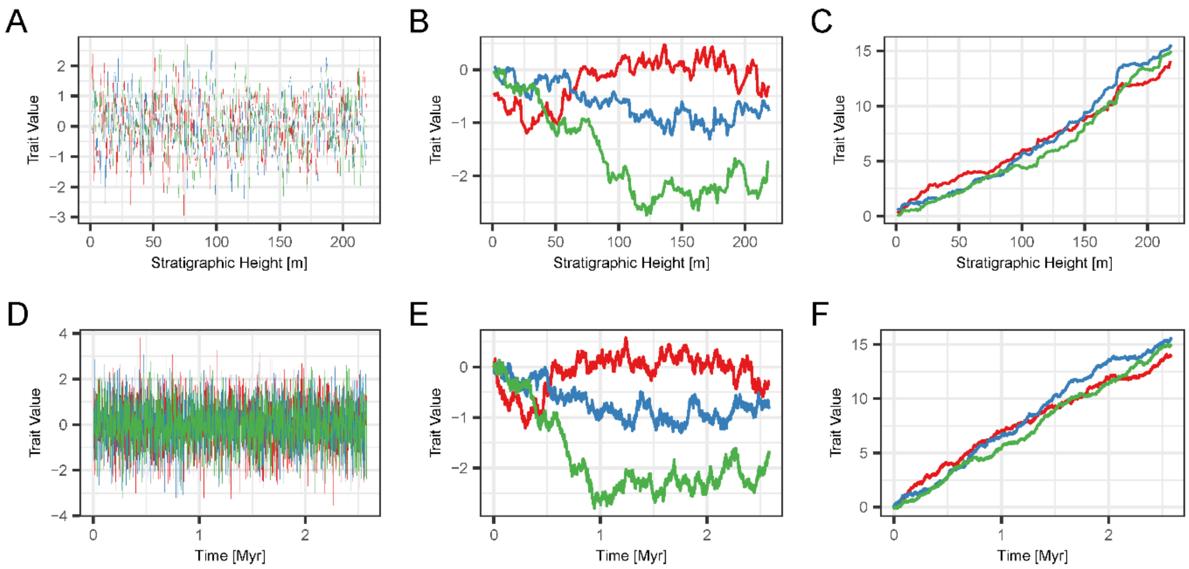


*Supplementary Figure 10: Differential preservation of different modes of evolution in scenario B, 8 km from shore. First row: preservation of three lineages evolving according to the stasis (A), Brownian motion (B), and Brownian drift (C) model. Second row: The corresponding true evolutionary history in the time domain.*

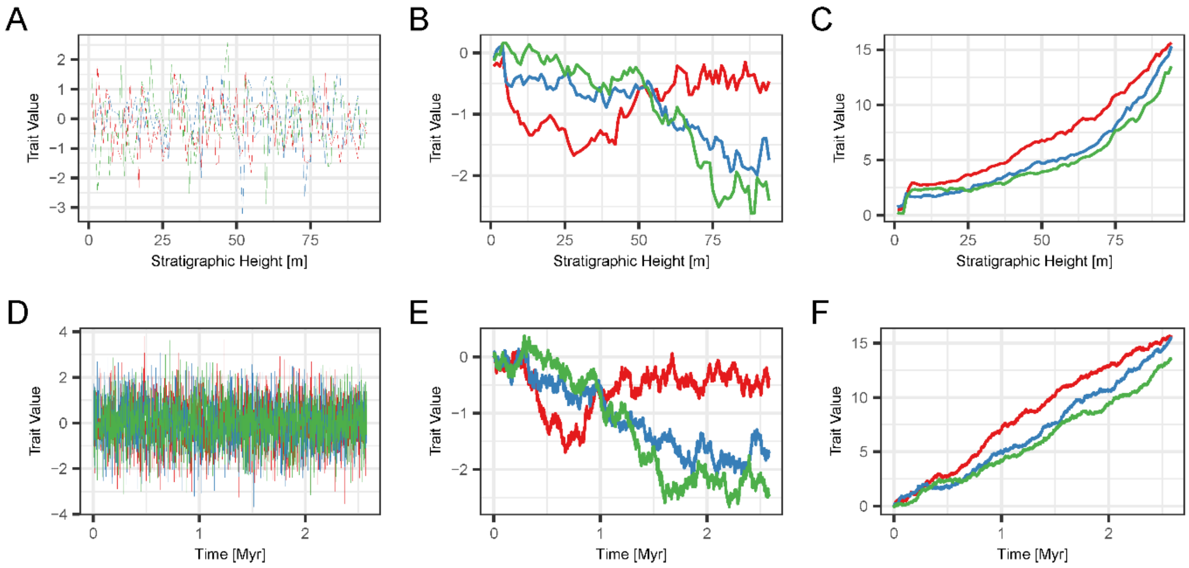


*Supplementary Figure 11: Differential preservation of different modes of evolution in scenario B, 10 km from shore. First row: preservation of three lineages evolving according to*

*the stasis (A), Brownian motion (B), and Brownian drift (C) model. Second row: The corresponding true evolutionary history in the time domain.*

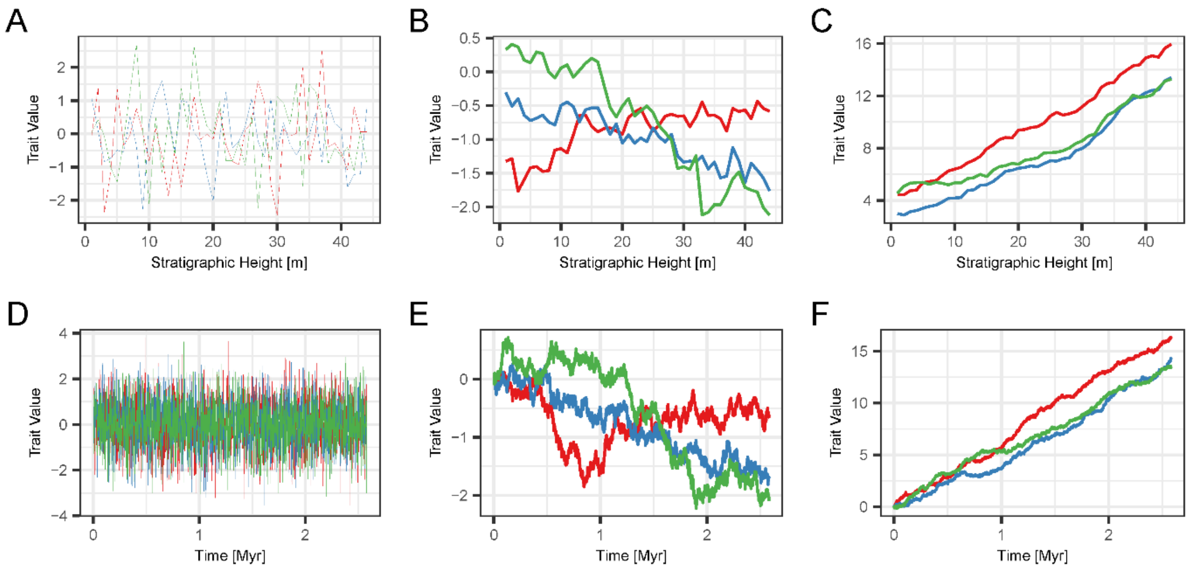


*Supplementary Figure 12: Differential preservation of different modes of evolution in scenario B, 12 km from shore. First row: preservation of three lineages evolving according to the stasis (A), Brownian motion (B), and Brownian drift (C) model. Second row: The corresponding true evolutionary history in the time domain.*

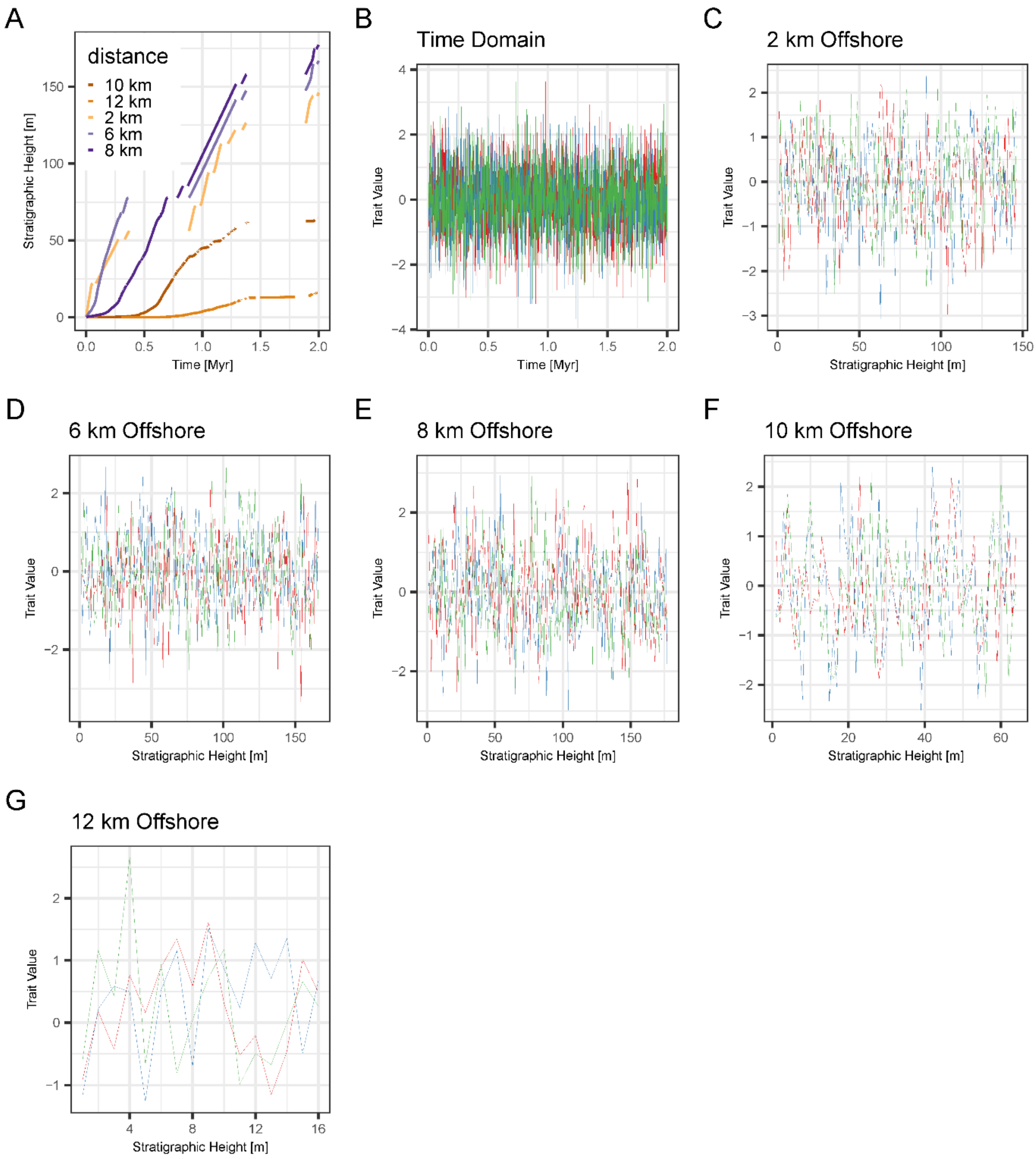


*Supplementary Figure 13: Preservation of stasis at different distances from shore in scenario A. (A): Age-depth models (B): three simulations of stasis in the time domain (C), (D), (E), (F), (G): preservation of the lineages from (B) in the stratigraphic domain at 2 km, 6 km, 8 km, 10 km, and 12 km offshore.*

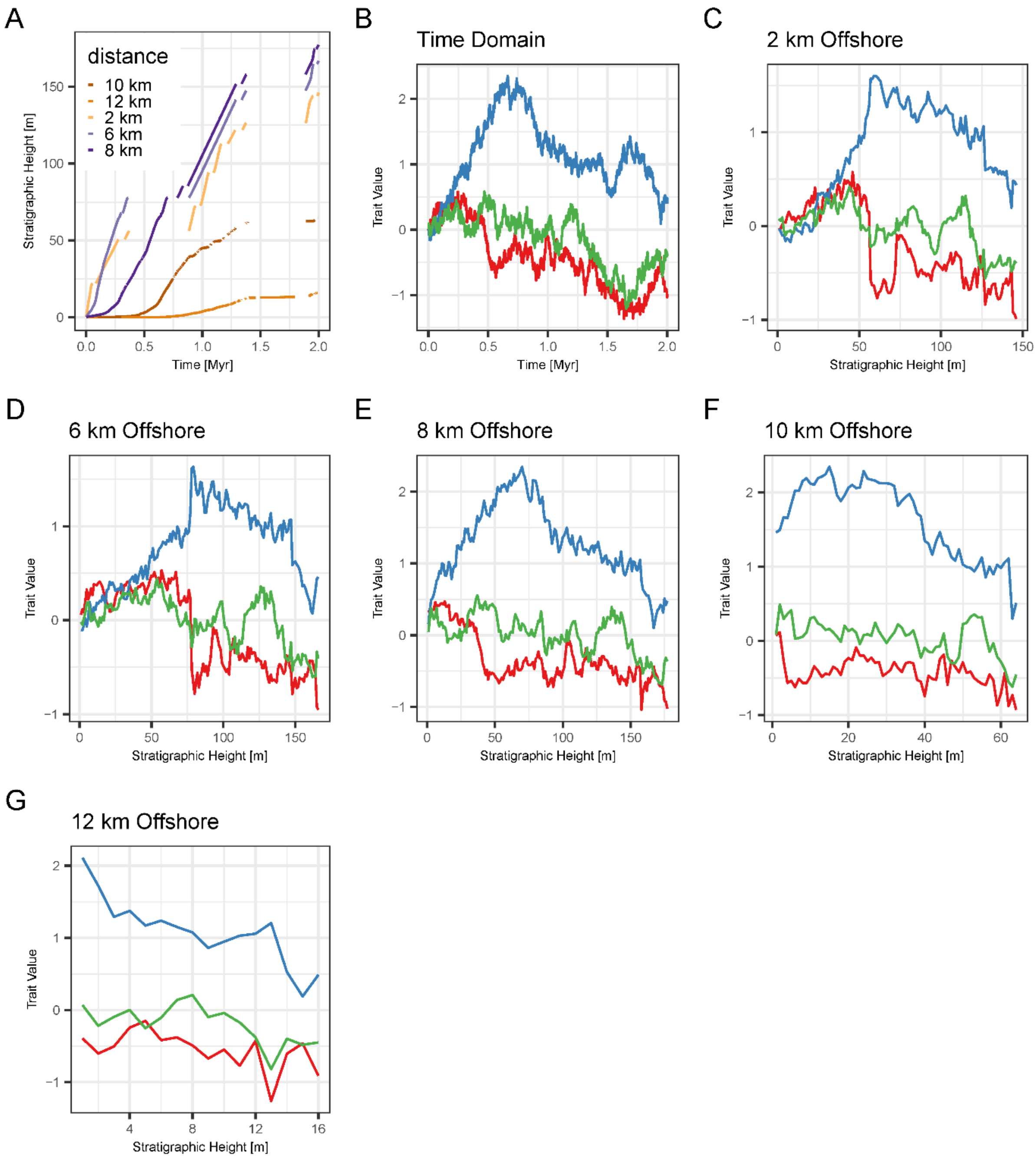


*Supplementary Figure 14: Preservation of Brownian motion at different distances from shore in scenario A. (A): Age-depth models (B): three simulations of Brownian motion in the time domain (C), (D), (E), (F), (G): preservation of the lineages from (B) in the stratigraphic domain at 2 km, 6 km, 8 km, 10 km, and 12 km offshore.*

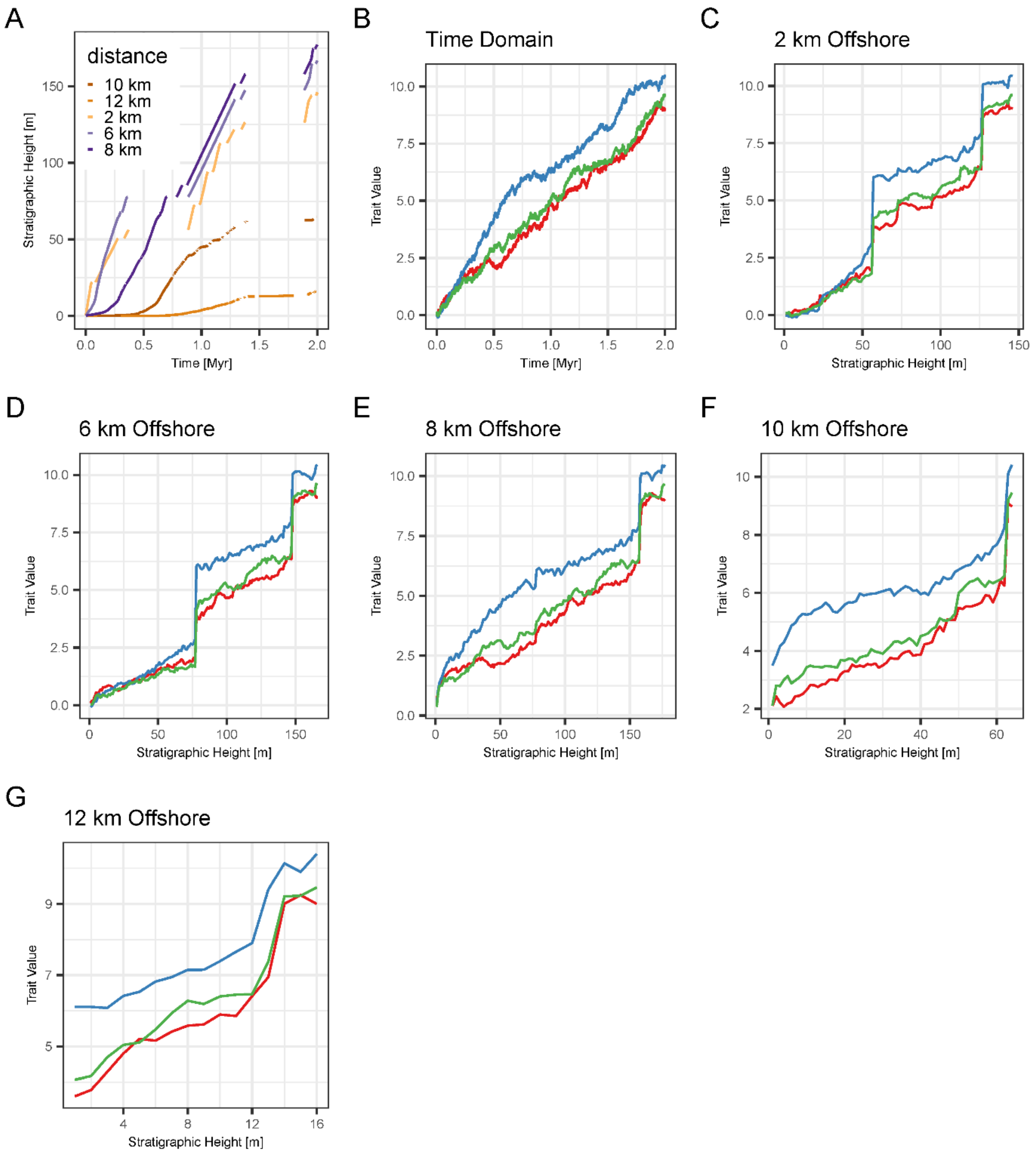


*Supplementary Figure 15: Preservation of Brownian drift at different distances from shore in scenario A. (A): Age-depth models (B): three simulations of Brownian drift in the time domain (C), (D), (E), (F), (G): preservation of the lineages from (B) in the stratigraphic domain at 2 km, 6 km, 8 km, 10 km, and 12 km offshore.*

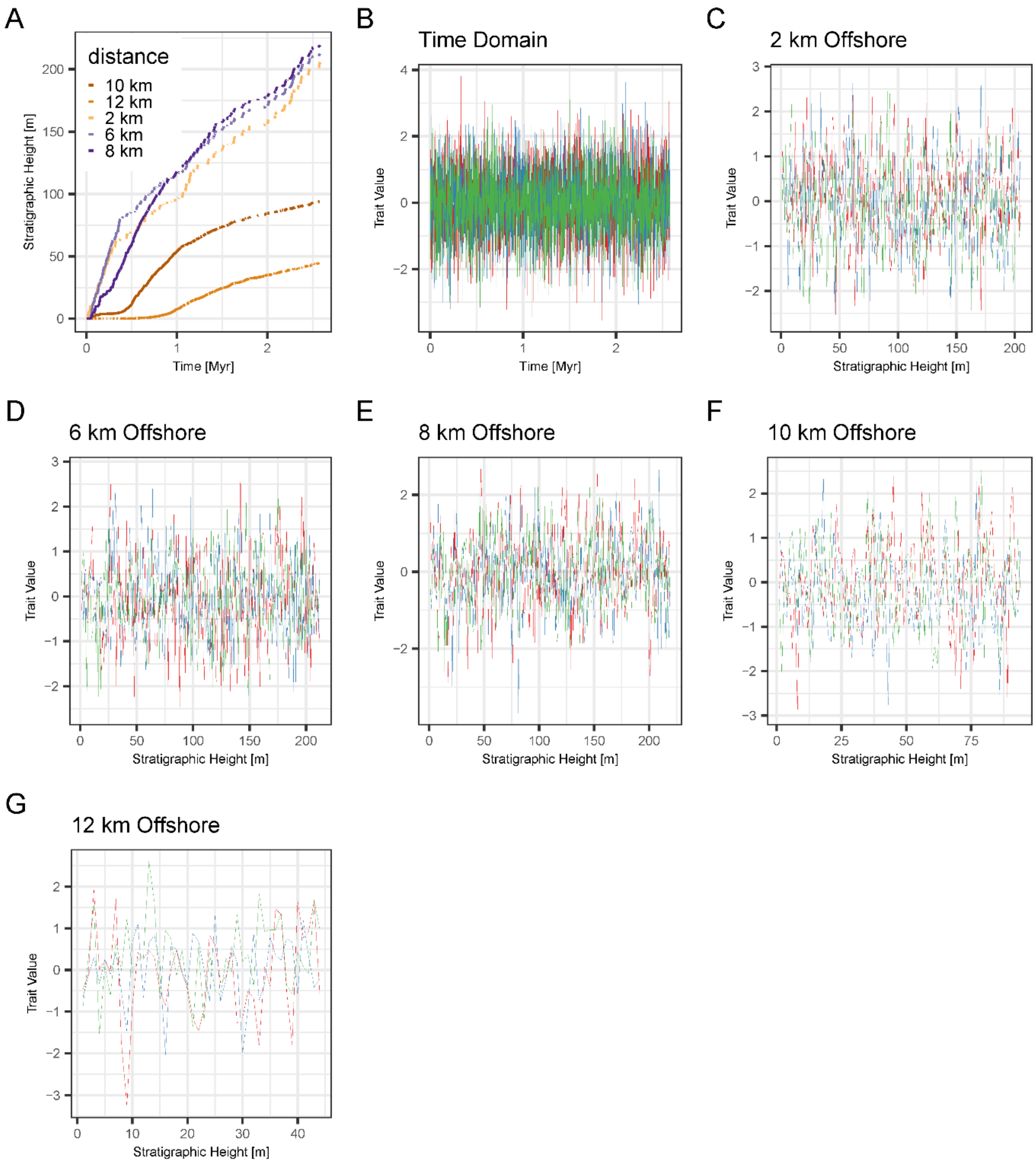


*Supplementary Figure 16: Preservation of stasis at different distances from shore in scenario B. (A): Age-depth models (B): three simulations of stasis in the time domain (C), (D), (E), (F), (G): preservation of the lineages from (B) in the stratigraphic domain at 2 km, 6 km, 8 km, 10 km, and 12 km offshore.*

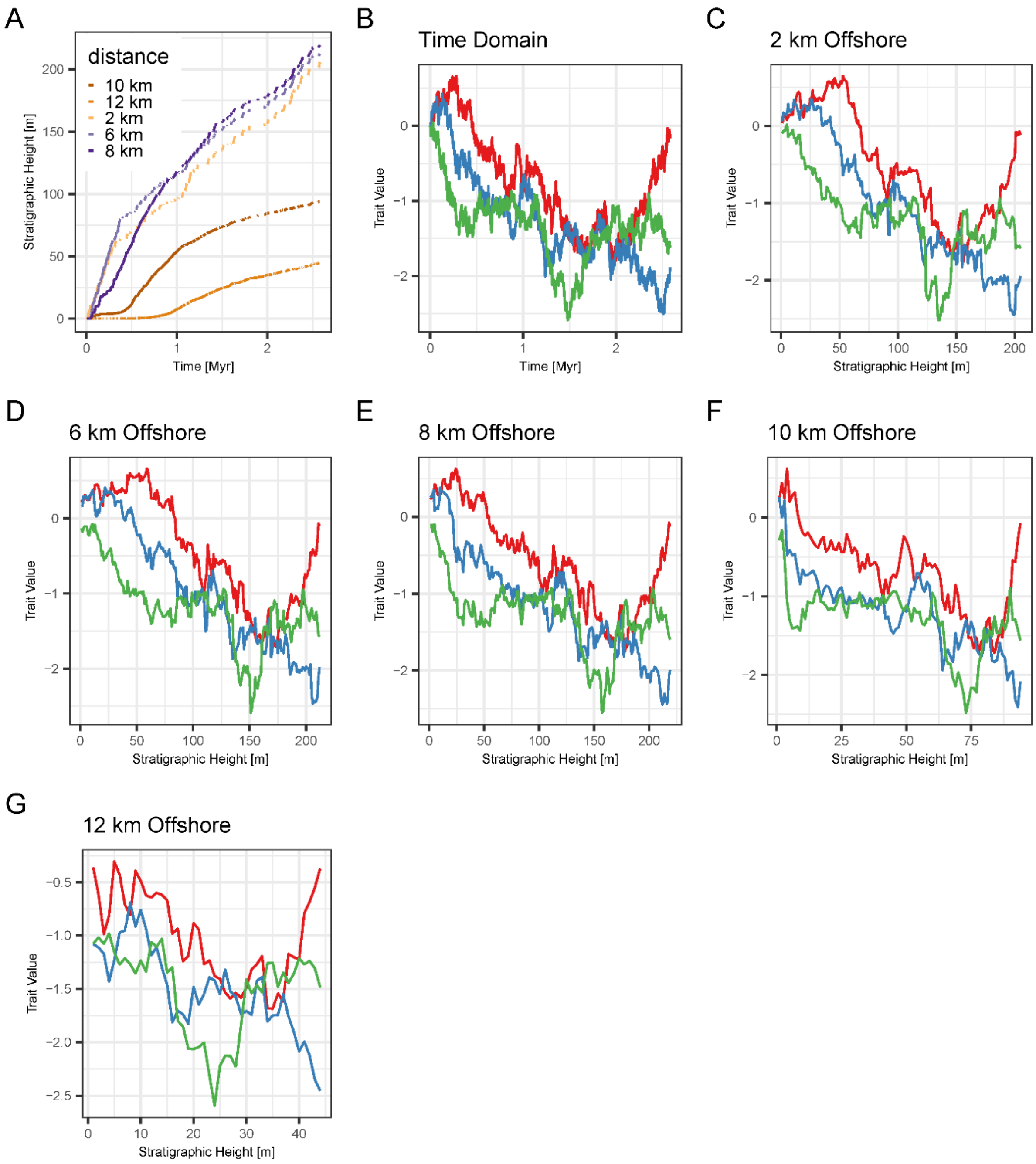


*Supplementary Figure 17: Preservation of Brownian motion at different distances from shore in scenario B. (A): Age-depth models (B): three simulations of Brownian motion in the time domain (C), (D), (E), (F), (G): preservation of the lineages from (B) in the stratigraphic domain at 2 km, 6 km, 8 km, 10 km, and 12 km offshore.*

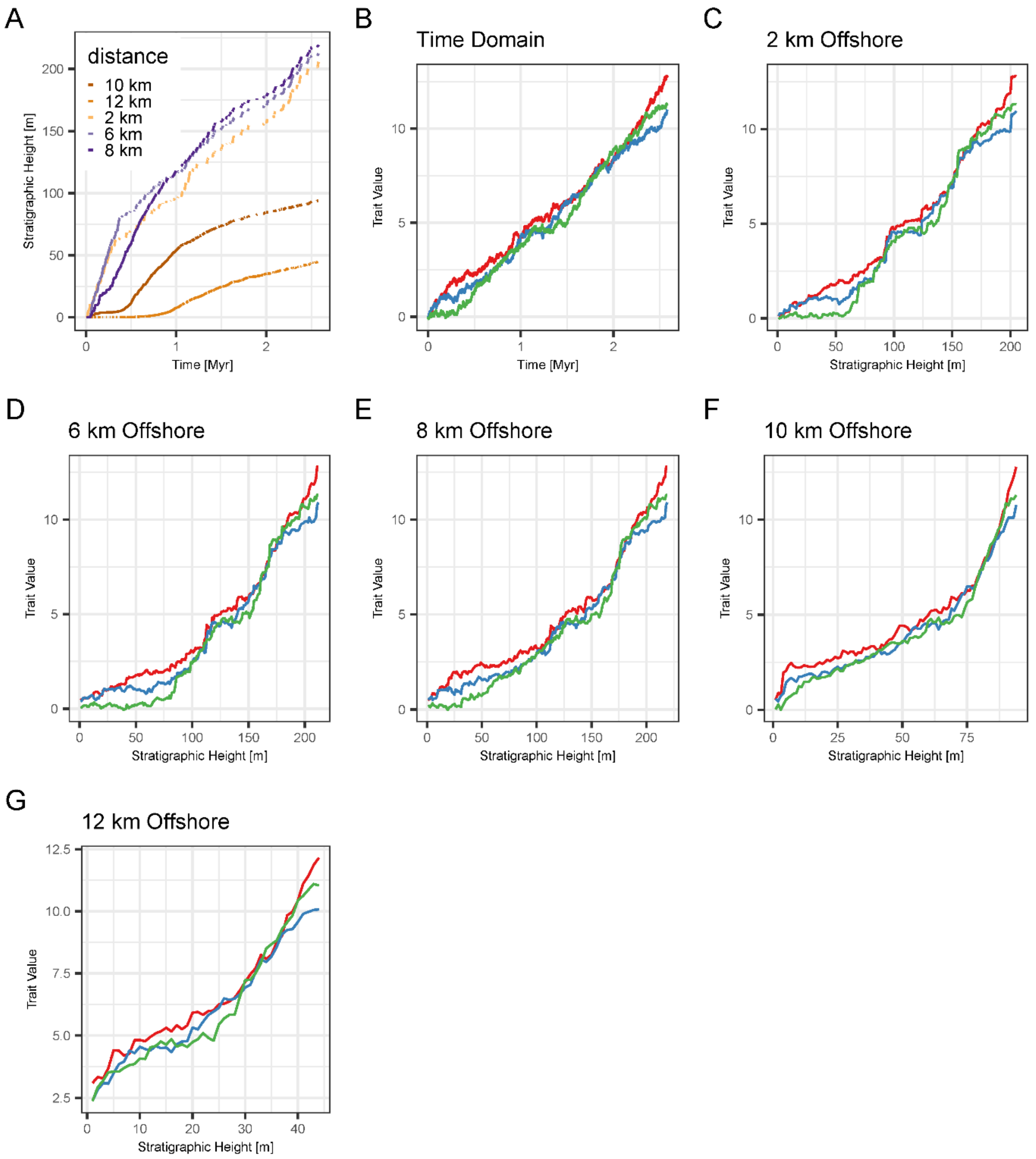


*Supplementary Figure 18: Preservation of Brownian drift at different distances from shore in scenario B. (A): Age-depth models (B): three simulations of Brownian drift in the time domain (C), (D), (E), (F), (G): preservation of the lineages from (B) in the stratigraphic domain at 2 km, 6 km, 8 km, 10 km, and 12 km offshore.*

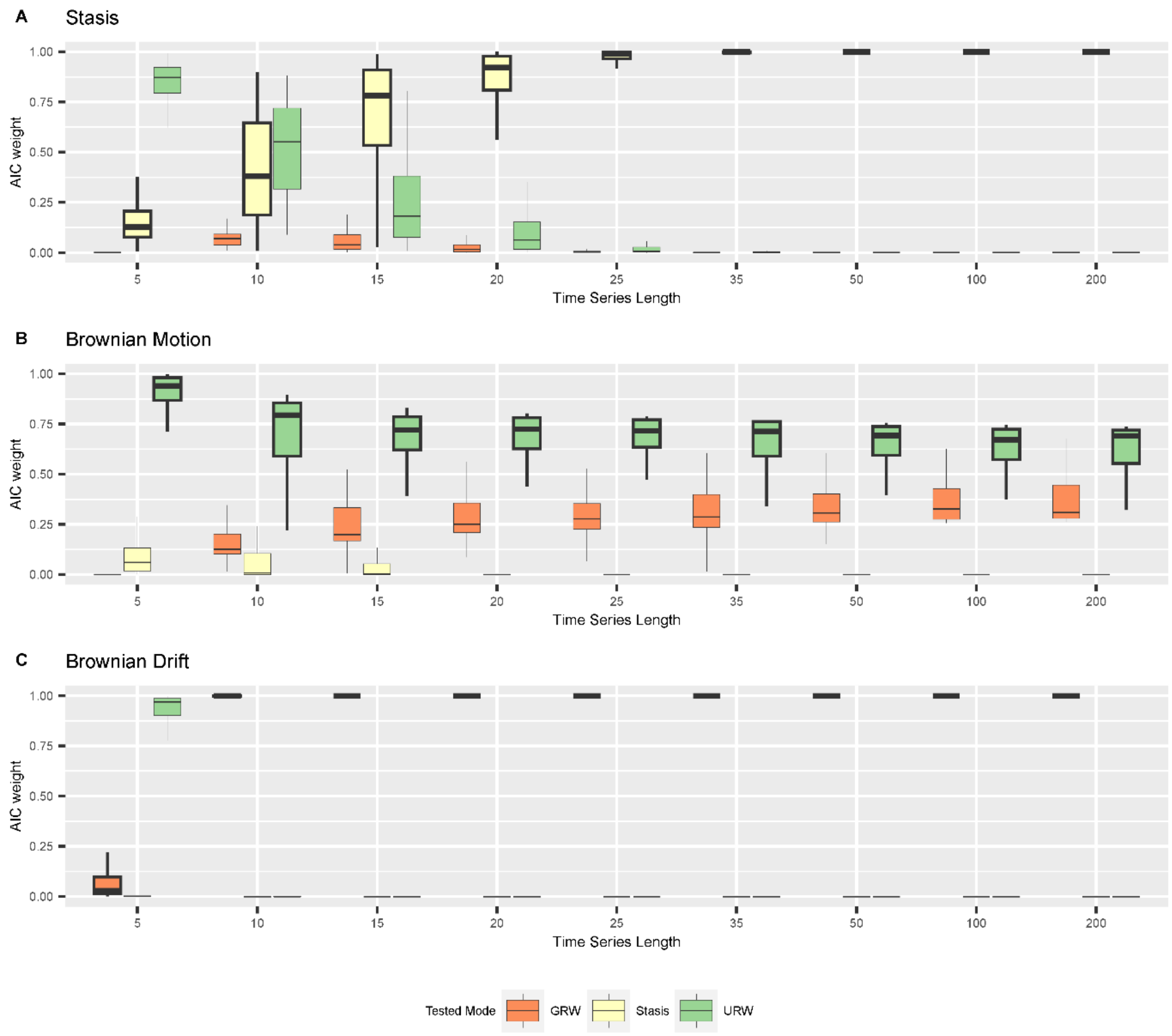


*Supplementary Figure19: AIC weights of modes of evolution in the time domain under simulations of (A) stasis; (B) Brownian motion; (C) Brownian drift as a function of the number of time series length. The sampled time interval is 2.58 Ma long, corresponding to the duration of scenario B.*

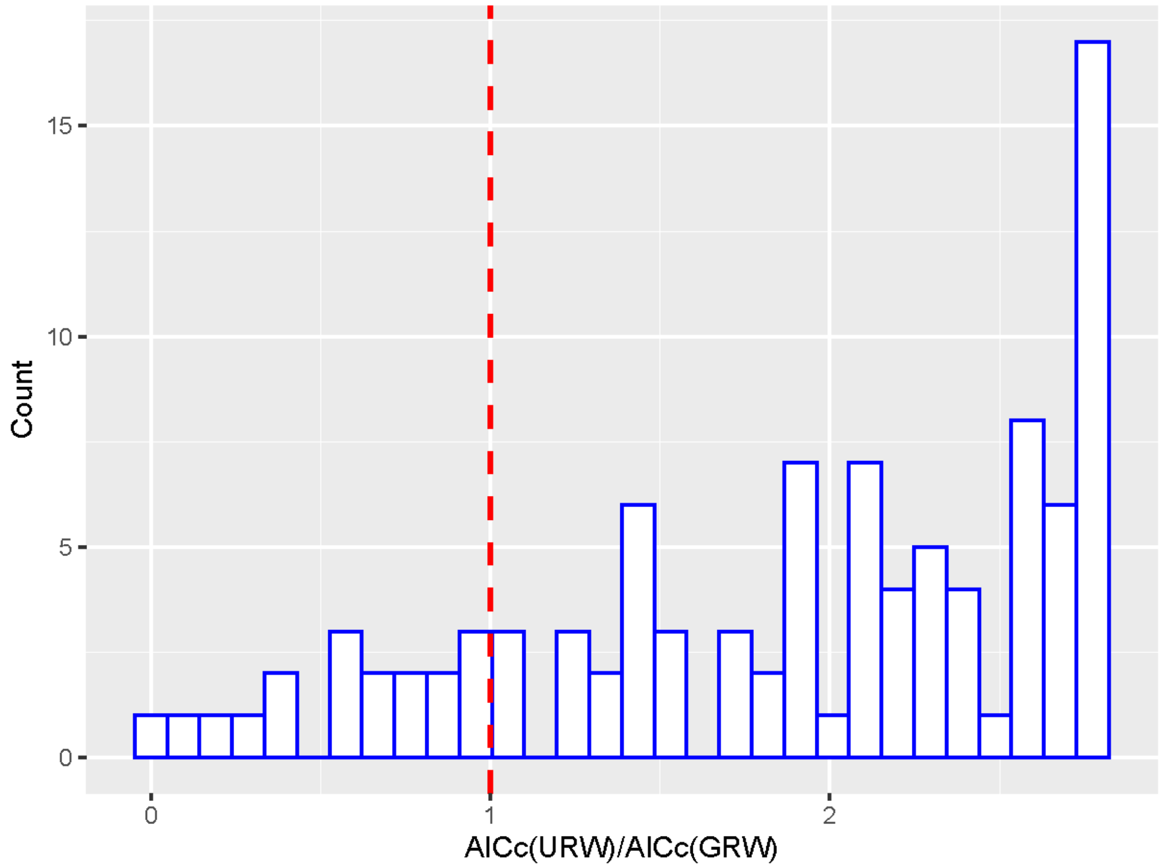


*Supplementary Figure 20: Evidence ratio (ratio of AICc weights) of undirected random walk (URW) and generalized random walk (GRW) under simulations of Brownian motion. An evidence ratio below 1 indicates that there is stronger support for the incorrect mode of evolution (GRW) than for the correct mode (URW).*
